# Sexual Dimorphism in VEMP peak to trough Latency

**DOI:** 10.1101/2023.04.14.536930

**Authors:** Max Gattie, Elena V. M. Lieven, Karolina Kluk

**Affiliations:** Manchester Centre for Audiology & Deafness (ManCAD), University of Manchester, Oxford Road, Manchester, M13 9PL, United Kingdom; Child Study Centre, The University of Manchester, Oxford Road, Manchester, M13 9PL, United Kingdom; The ESRC International Centre for Language and Communicative Development (LuCiD), The University of Manchester, Oxford Road, Manchester, M13 9PL, United Kingdom

## Abstract

The cervical vestibular-evoked myogenic potential (VEMP) was assessed in 24 women and 24 men having a mean age of 19.5 years (SD 0.7). Whilst there was no group difference in VEMP peak to trough (p1-n1) amplitude, VEMP p1-n1 latency was found to be shorter for women than for men by 2.4 ms (95% CI [–0.9, –3.9], chi squared (1) 9.6, p = 0.0020). This equates to 21% of the mean 11.4 ms VEMP p1-n1 latency across women and men. It is a reversal of findings in several prior studies, which are reviewed here. Statistical modelling based on the current study suggests some prior studies were underpowered to detect a sex difference in VEMP latency. Possible causes for sex difference in VEMPs are discussed. Candidate explanations include head resonance, superposition of motor unit action potentials and influence of sex hormones. These explanations are not mutually exclusive, and multiple factors may contribute to difference in VEMP measurement between women and men. This study used a methodology developed in Gattie et al. (2021), which addresses sound exposure concerns with the high amplitude air conducted stimuli necessary to evoke a VEMP response. It is suggested that body conducted stimuli may be preferable for VEMP testing in which ear-specific information is not required.

## 1 Introduction

Women have been identified more frequently than men in a variety of diagnoses related to the vestibular system (Smith et al., 2019). In a review of healthcare data representing approximately 86% of the German population (38,377,351 women and 31,938,568 men), women were found to be more frequently affected than men by dizziness and vertigo (Hülse et al., 2019). There was moreover a finding of higher prevalence of vestibular-related diagnoses between the ages of 15 and 74 years, including Benign Paroxysmal Positional Vertigo, Vestibular Vertigo, Vestibular Neuritis, Vestibular Migraine, mal de debarquement syndrome, Unspecified Peripheral Vestibular Dizziness and Motion Sickness. Mal de debarquement syndrome, in which the sensation of regular motion persists long after exposure to regular motion has ceased, has long been recognised to have higher prevalence in women than men (Hain & Cherchi, 2016; Cha et al., 2018).

Following its initial description in the 1990s (Colebatch & Halmagyi, 1992) the cervical vestibular-evoked myogenic potential (VEMP) has become accepted as a reliable test of vestibular function. It measures a short inhibition of tonic activity in the sternocleidomastoid muscle (SCM) (Corneil & Camp, 2018; Rosengren & Colebatch, 2018) and can be thought of as a short latency fragment of the vestibulo-collic reflex (Forbes et al., 2018). As illustrated in figure 1, the VEMP is thought to correspond to a reflex arc involving the vestibular periphery, the VIII cranial nerve, vestibular nuclei, the medial vestibulospinal tract, the XI cranial nerve and the SCM. The exact trajectory through vestibular nuclei is subject to ongoing investigation (Forbes et al., 2013).

**Figure 1.**
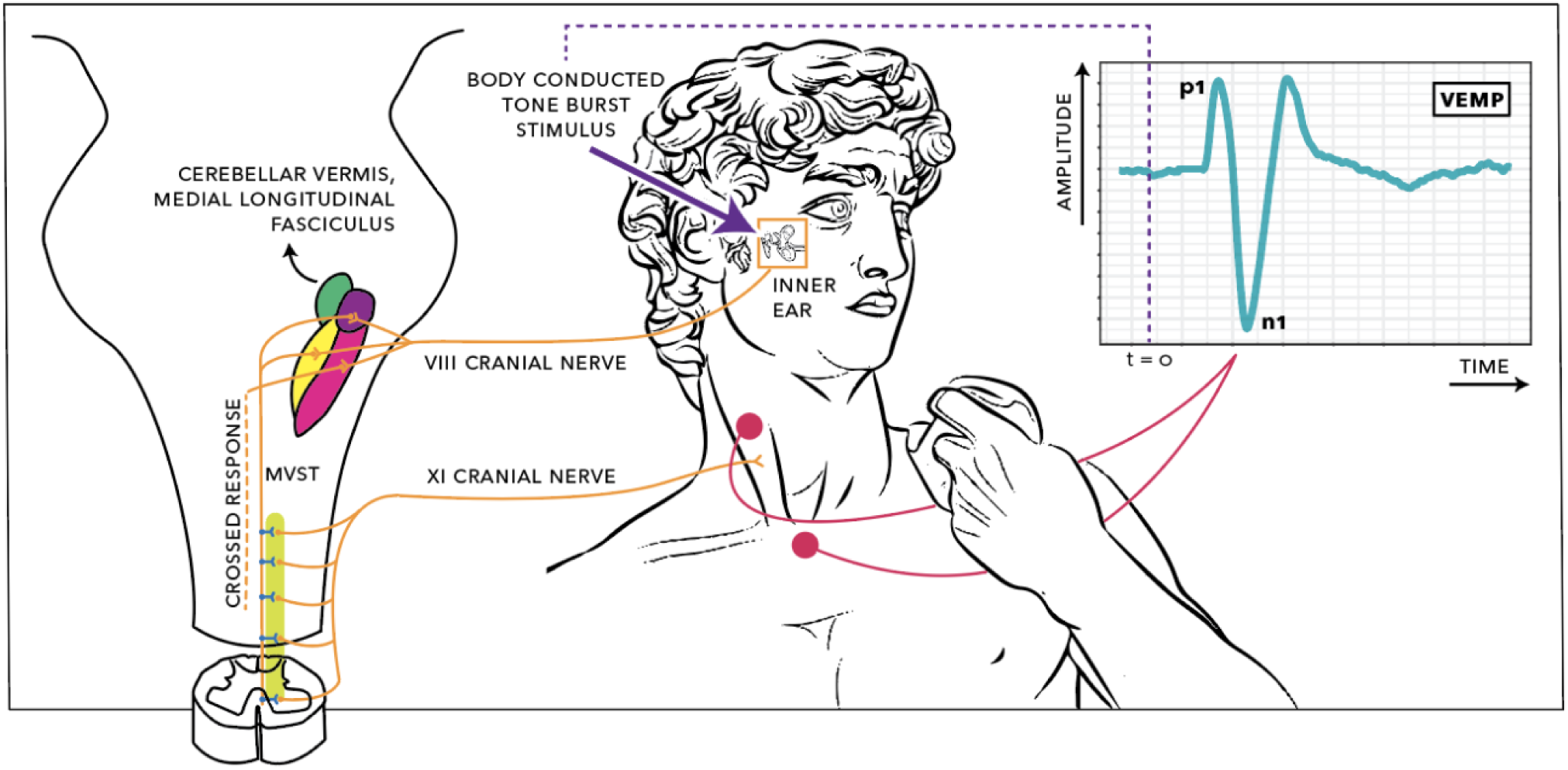
Electrophysiological recording arrangement for cervical vestibular-evoked myogenic potentials (VEMPs). The stimulus is a transient delivered through vibration, or high amplitude air conduction. Energy from the transient deflects hair cells in the vestibular system, creating an electrical impulse along the VIII cranial nerve. These vestibular afferents synapse in the vestibular nucleus then descend along the medial vestibulospinal tract (MVST). The MVST connects to the XI cranial nerve, which branches to synapse on motoneurons innervating the ipsilateral sternocleidomastoid. Activity following delivery of a transient results in a brief inhibition of tonic activity in the muscle spindles of the sternocleidomastoid. This inhibitory potential can be recorded between electrodes on the sternum and the belly of the sternocleidomastoid, creating a wave form with a characteristic peak (p1) and trough (n1) at approximately 13 ms and 23 ms after stimulus onset. The vestibulo-collic reflex also includes crossed and ascending components, not shown in detail. Based on Kim et al. (2010), Oh et al. (2016), and Colebatch et al. (2016). Creative Commons CC BY-NC-SA 4.0.

The VEMP is a large potential with a characteristic peak (p1) and trough (n1). It has been described as a superposition of motor unit action potentials (Wit & Kingma, 2006; Lütkenhöner, 2019). Wei et al. (2013) measured tuning curves with nine frequencies between 125 and 4000 Hz, and suggested that VEMP could be modelled as linear summation of two mass spring systems, with resonance frequencies at approximately 300 Hz and 1000 Hz. Rosengren et al. (2016) used concentric needle electrodes to measure VEMP in human participants. Their suggestion was that p1 behaves as a travelling wave, whereas n1 is a combination of a trailing dipole following the propagating inhibition, a standing wave generated when the inhibition reaches the end of the muscle, and a small rebound in firing following inhibition.

Table 1 shows studies which have compared measures of VEMP p1 and n1 amplitude and latency between women and men. No consistent sex difference is apparent.

**Table 1.**
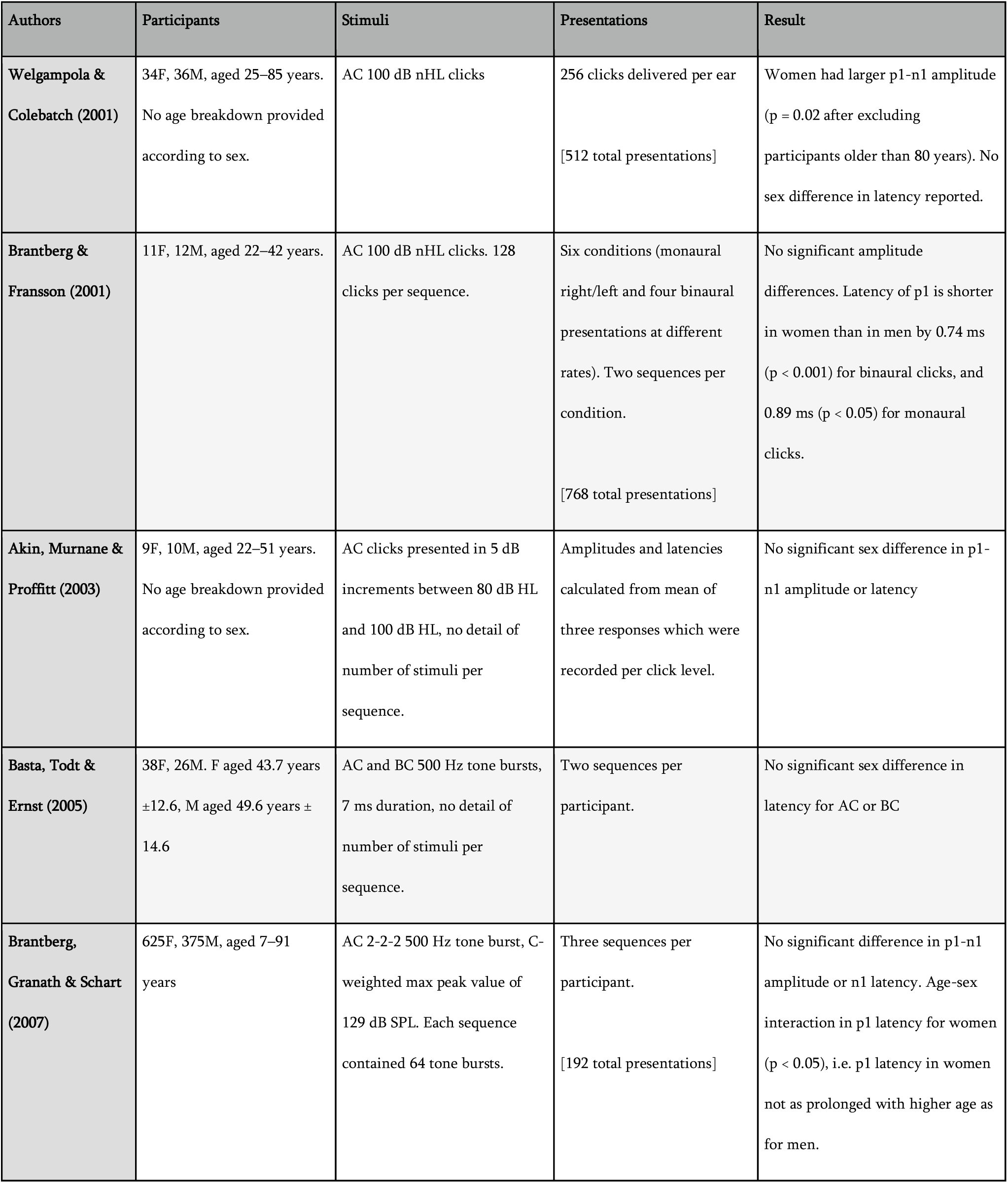
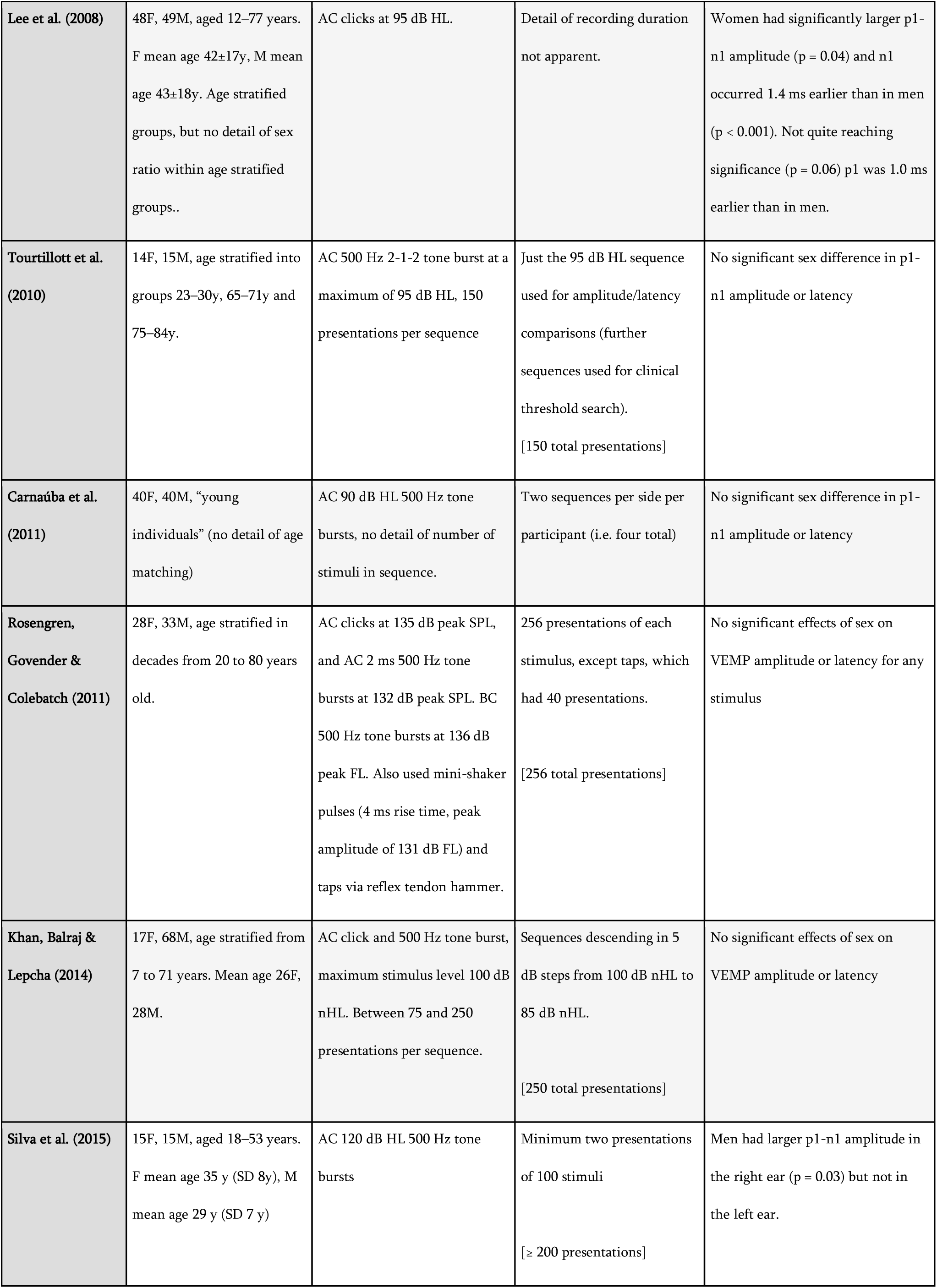
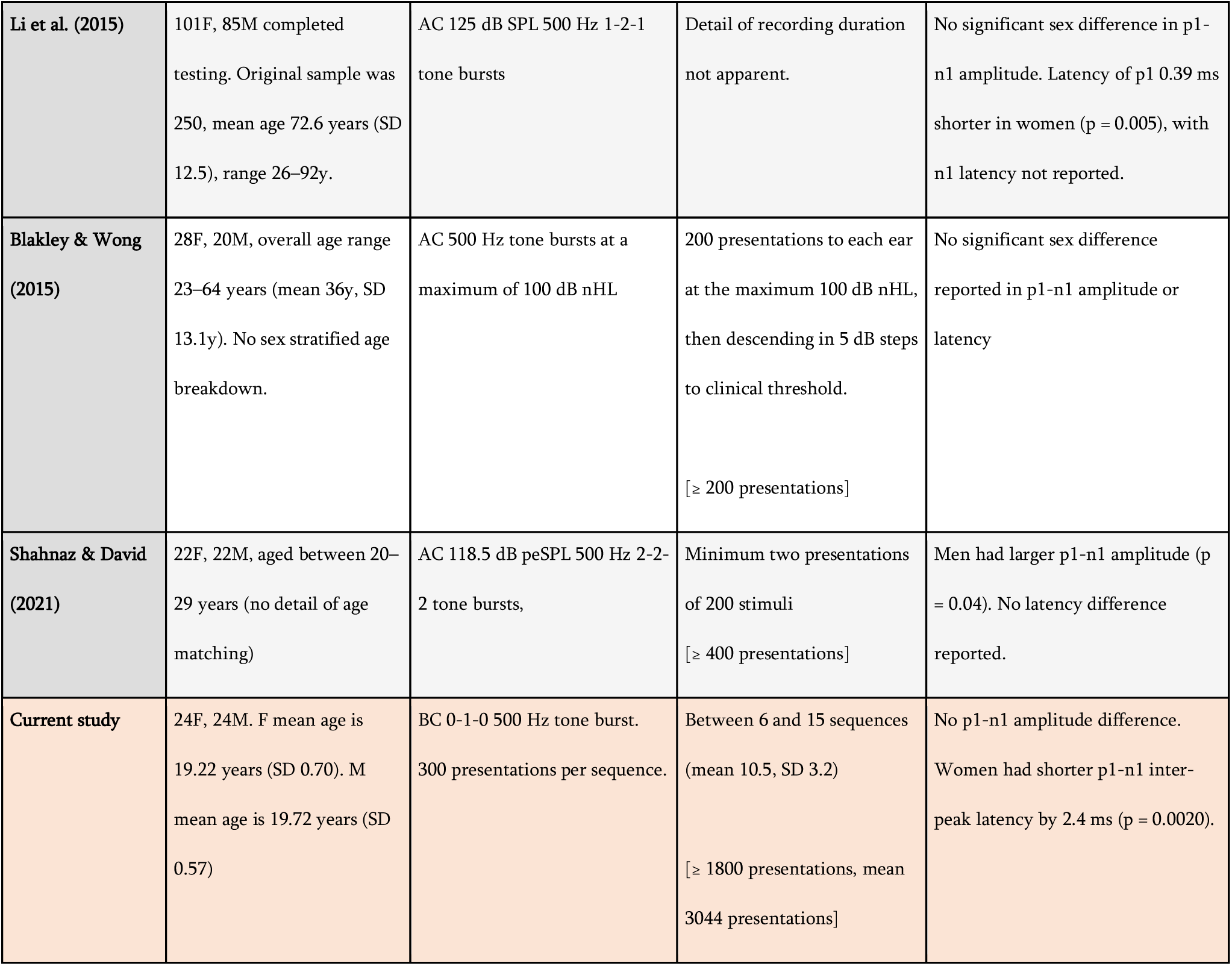
Studies comparing cervical vestibular-evoked myogentic potential (VEMP) latencies in women (F) and men (M). Stimuli were air-conducted (AC) in all but the current study, which used bone-conducted (BC) stimuli. The units used to report stimuli are inconsistent. See section 2.2 for discussion of how units might be compared.

The current study assessed VEMP using a methodology developed for a pre-registered project (Gattie et al., 2019) assessing differences between groups who do and do not stutter (Gattie et al., 2021). The study appraised VEMP amplitude and latency, using peak to trough measures with SCM tension controlled using biofeedback and a custom head bar apparatus, and analysis via linear mixed-effects regression modelling. The methodology was developed with the aim of ensuring adequate statistical power to test the pre-registered hypotheses. This was achieved by using body-conducted (BC) stimuli throughout. The incentive to use BC stimuli followed from consideration of safe sound levels. Sudden bilateral hearing loss has been reported following VEMP testing (Mattingly et al., 2015; Asakura & Kamogashira, 2021), raising concerns about the very high air-conducted (AC) stimulus levels required to evoke a VEMP response.

BC stimuli elicit a VEMP response at appreciably lower levels than are necessary when using AC stimuli (McNerney & Burkard, 2011). Thus, use of BC stimuli can enable collection of substantially more data than if AC stimuli had been used, thereby increasing statistical power. Data presented here are part of normative data collection with a student population. Power analysis, including appraisal of participant count, is described in detail in section 4.1.

## 2 Materials and Methods

### 2.1 Participants

Participants were students, of whom 47 were undergraduates participating in exchange for course credit, and one was an A-level student on placement. Equal numbers of women and men participated. Women were aged between 16.6 and 21.1 years (mean 19.22, SD 0.70) and men were aged between 18.6 and 20.7 years (mean 19.72, SD 0.57). Otoscopy, tympanometry and pure tone audiometry were performed for all participants following procedures described by the British Society of Audiology (BSA 2014, 2018, 2022), with results within the normal range. All participants gave written informed consent according to the Declaration of Helsinki. The University of Manchester Ethics Committee approved the study.

### 2.2 Experimental Arrangement

VEMP was assessed with an Eclipse EP25 system (Interacoustics AS, Assens, Denmark). Skin was prepared with NuPrep® (Weaver and Company, CO, United States) prior to electrode attachment with Ten20® conductive paste (Weaver and Company, CO, United States). Non-metallic silver chloride disposable electrodes were used (type M0835, Biosense Medical, Essex, United Kingdom). Electrode impedance was below 3 kΩ. The montage included an active electrode on the right hand side SCM, and reference and ground electrodes on the upper sternum and nasion respectively.

Stimuli were 500 Hz tone bursts with rectangular windowing, created by the Eclipse. The frequency of 500 Hz has been shown as optimal for VEMP testing (Rosengren et al., 2010; Papathanasiou et al., 2014). BC and AC stimuli were both tested, although only results with AC stimuli are reported here. Characteristics of BC stimuli were matched those of AC stimuli to facilitate comparison between BC and AC VEMPs. Following the considerations around safe sound levels described in section 1, the shortest possible AC stimuli were used. These were 0-1-0 tone bursts with a rise/fall time of zero and a plateau of 2 ms, for an overall characteristic between a tone burst and a click (Laukli & Burkard, 2015). BC stimuli were delivered to the mastoid bone behind the right ear at a rate of 5.1 every second, using a B81 bone conductor (Radioear, MN, United States).

The studies in table 1 report stimulus levels inconsistently. This report presents BC stimulus levels using hearing level (HL) equivalent units, rather than the force level units which are typically used to report vibratory stimuli. Doing so facilitates comparison with studies in table 1. However, it should be emphasised that the dB HL scale used in this report is dissimilar to the HL scales used for pure tone audiometry (e.g. dB HL re: ANSI S3.6-1996 or dB HL re: ISO 389-1:2017) and is also dissimilar from the dB nHL and eHL scales used in ABR testing (NHSP Clinical Group, 2013). This was unavoidable due to absence of a reference equivalent threshold for the transducer and stimulus used, and reflects ongong debate over standardisation of calibration procedures for acoustic transients (Burkard, 2006; Lightfoot et al., 2007; Laukli & Burkard, 2015). The procedure for calibration of the bone conductor used a Model 4930 artificial mastoid and 2250 Investigator (Brüel and Kjaer, Naerum, Denmark), and Agilent 54621A 2-Channel Oscilloscope (Keysight, CA, United States). The artificial mastoid had a reference equivalent threshold force level re 1 μN of 40.2 dB for 500 Hz. This reference equivalence was used with a correction factor, provided by Interacoustics, of 69.5 dB for peSPL to nHL conversion of a 2-2-2 500 Hz tone burst; however, and as already noted, this was applied to a 0-1-0 500 Hz tone burst. Based on normative data (e.g. Beattie & Rochverger, 2001; Gorga et al., 2006), inaccuracy introduced by doing so is unlikely to be greater than 5 dB. Any inaccuracy in the threshold reference will moreover not have affected between group comparisons, since both groups were equally affected. This calibration method enabled the amplitude of the bone-conducted 0-1-0 500 Hz stimulus to be reported in dB HL units. Examination of the output using the artificial mastoid and an oscilloscope showed clipping of the trace when amplitudes exceeded 40 dB HL. Stimuli were accordingly set so they would not exceed 40 dB HL.

The Eclipse was used to amplify, record and filter the electromyography (EMG) signal using the Interacoustics research license. A digital FIR filter of 102nd order was set to low pass at 1500 Hz, and an analog Butterworth filter of 1st order at 6 dB per octave was set to high pass at 10 Hz. Sampling by the Eclipse was at the highest resolution available for VEMP protocols, which was 3 kHz. The 3 kHz sampling rate meant that measurements of VEMP p1-n1 latency were grouped into steps of 0.33 ms. Such grouping is visible in figures 11 and 12.

### 2.3 Procedure

Participants sat with their foreheads resting against a padded bar. The apparatus was specifically constructed for the experiment (figure 2). Participants were asked to push their heads on the padded bar and try to maintain an EMG biofeedback target as close as possible to 50 µV root mean square (RMS). If background EMG fell lower than 50 µV RMS, stimuli stopped playing and participants were asked to push harder. Participants were instructed not to push harder than necessary to keep the stimuli playing, and would rarely attempt to do so. The importance of maintaining a constant background EMG was explained. Background EMG was monitored by the experimenter throughout testing.

**Figure 2:**
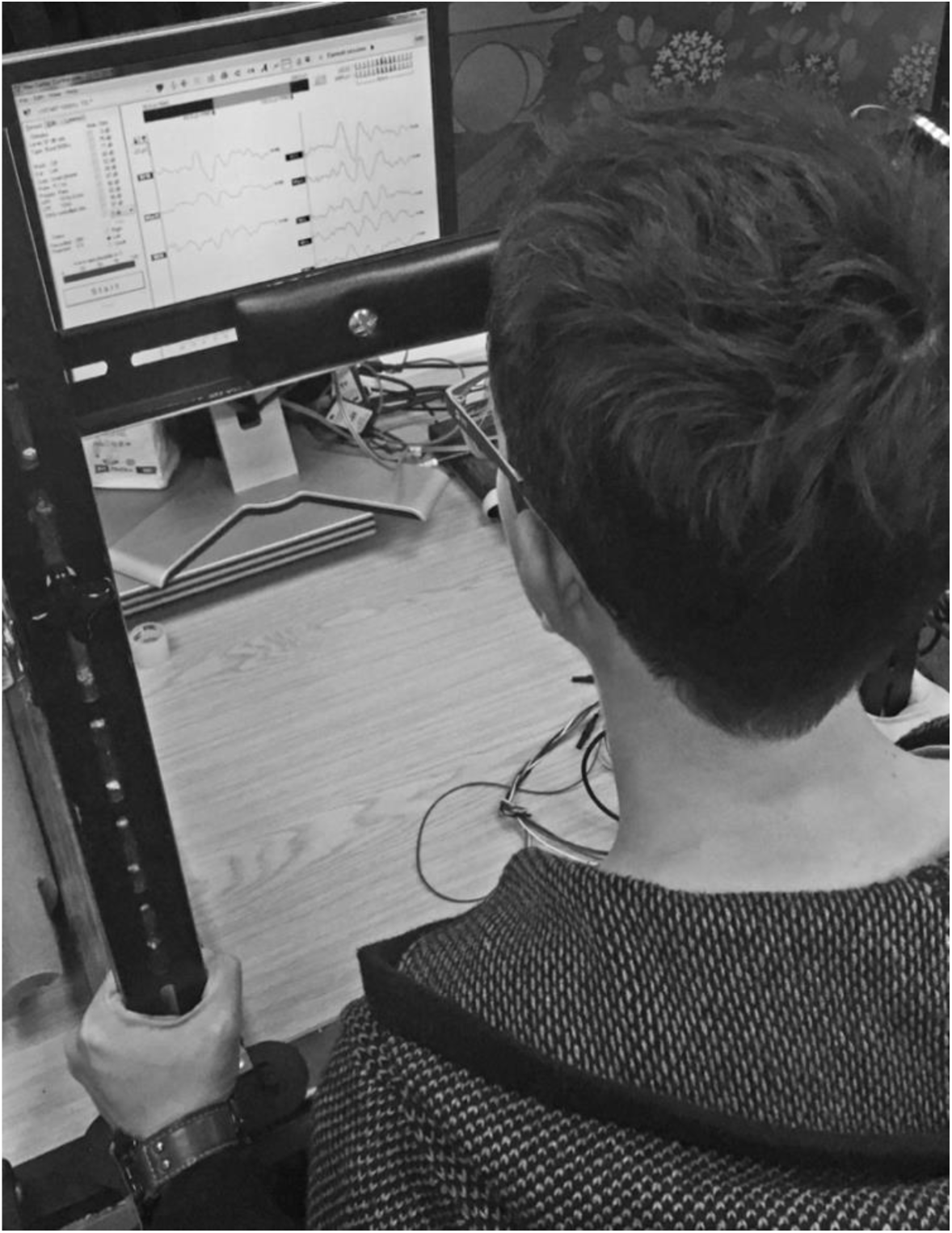
Participants were asked to push against a padded bar with their foreheads, maintaining tension in the sternocleidomastoid tension as close as possible to 50 µV root mean square throughout testing. The Eclipse clinical software provided biofeedback helping participants to monitor sternocleidomastoid tension.

Recordings followed the procedure recommended by Interacoustics for VEMPs. Epochs with peak or trough amplitudes having a magnitude larger than ± 800 µV were rejected. The Eclipse software compensated for rejected epochs, so that for every stimulus level tested an averaged response to exactly 300 epochs was recorded. These averages of 300 epochs will henceforth be referred to as “sequences”. Recording of the initial sequence was at a stimulus level of 40 dB HL, and further sequences were recorded with the stimulus level decreased in steps of 2 dB until either a presentation at 34 dB HL, or until the averaged VEMP trace which summarised the sequence was comparable to background noise, whichever came soonest. The experimenter compared the averaged VEMP trace to background noise using the EP25 clinical software. A second series of recordings started at 39 dB HL, with stimulus level decreased in 2 dB steps until either a presentation at 35 dB HL or until the averaged VEMP trace which summarised the sequence was comparable to background noise, whichever came soonest. The plan for data collection was described to participants, who were able to watch their averaged VEMP traces calculated in real time by the EP25 software. If participants were willing (e.g. in the event of no subsequent appointment) and they had shown a response at 34 dB HL, further sequences at stimulus levels below 34 dB HL were recorded. To complete the session, a repeat recording was made using the maximum 40 dB HL stimulus level.

### 2.4 Data processing

Custom scripts were written in MATLAB 2019a (The MathWorks, Inc., Natick, MA) and used to process raw data. Normalisation was carried out for each participant by transforming response amplitudes into a dimensionless ratio. This worked by extracting a pre-stimulus interval of 18 ms from a mean of the EMG waveforms from the first six sequences of 300 presentations recorded for each participant (i.e. the extract was a pre-stimulus mean of the first 1800 presentations recorded). A root mean square (RMS) based on this pre-stimulus mean was then assigned as a background EMG tension for each participant. Finally, each sequence was normalised by dividing it by the background EMG tension for its participant.

The normalisation procedure described was in principle not necessary, since use of the head bar limited variation in background EMG tension according to the 50 µV biofeedback target. However, the normalisation increased accuracy by adjusting for any small per participant variation in EMG tension. Normalisation used the maximum 1800 presentations available for every participant, and thus minimised random noise in the pre-stimulus RMS background EMG tension. This procedure was considered preferable to alternatives such as per sequence normalisation based on pre-stimulus RMS for each sequence of 300 presentations. Per sequence normalisation could have introduced noise because random fluctuation in pre-stimulus RMS per sequence (i.e. random in addition to any genuine change in sternocleidomastoid tension) would have randomly affected VEMP amplitudes on a per sequence basis, and thus affected within participant comparisons. Comparison of data between participants used linear mixed-effects regression analysis, and this in turn relied upon accurately measuring VEMP amplitude growth with stimulus level for each participant. Thus, preserving within participant comparisons as accurately as possible was optimal for linear mixed-effects regression analysis. The normalisation procedures used in this study preserved within participant comparisons with the same accuracy as that available from the raw data.

Figure 3 compares VEMP grand averages for women and men at the maximum 40 dB HL stimulus level. Peaks per sequence per participant were identified with the “findpeaks” algorithm in the MATLAB Signal Processing Toolbox. Troughs were identified by applying “findpeaks” to inverted waveforms. To begin with, peaks and troughs were identified for the initial 40 dB HL sequence per participant. This was done by first identifying all of the troughs in the 40 dB HL sequence, and then identifying as n1 the most prominent trough between 15–37 ms (with prominence defined as per the “findpeaks” algorithm). Following that, peaks were identified across the entire 40 dB HL trace. Peaks were discarded if they occurred earlier than 5 ms or later than n1. The peaks surviving this process were ranked. Firstly, the three peaks with greatest prominence were awarded 5, 4 and 3 points in order of prominence. Secondly, those three most prominent peaks were weighted according to their prominence relative to the most prominent peak: 3 points for greater than or equal to two thirds; 2 points for greater than one third and less than two thirds; and 1 point otherwise. Thirdly, the five peaks having latencies with the smallest time differences from n1 were given points from 5 to 1 in a hierarchy where higher points were awarded for a smaller time difference. Finally, all of the points were added together. The peak which had the greatest number of points was identified as p1. In the event of a tie, the chosen peak had the smallest time difference from n1.

**Figure 3:**
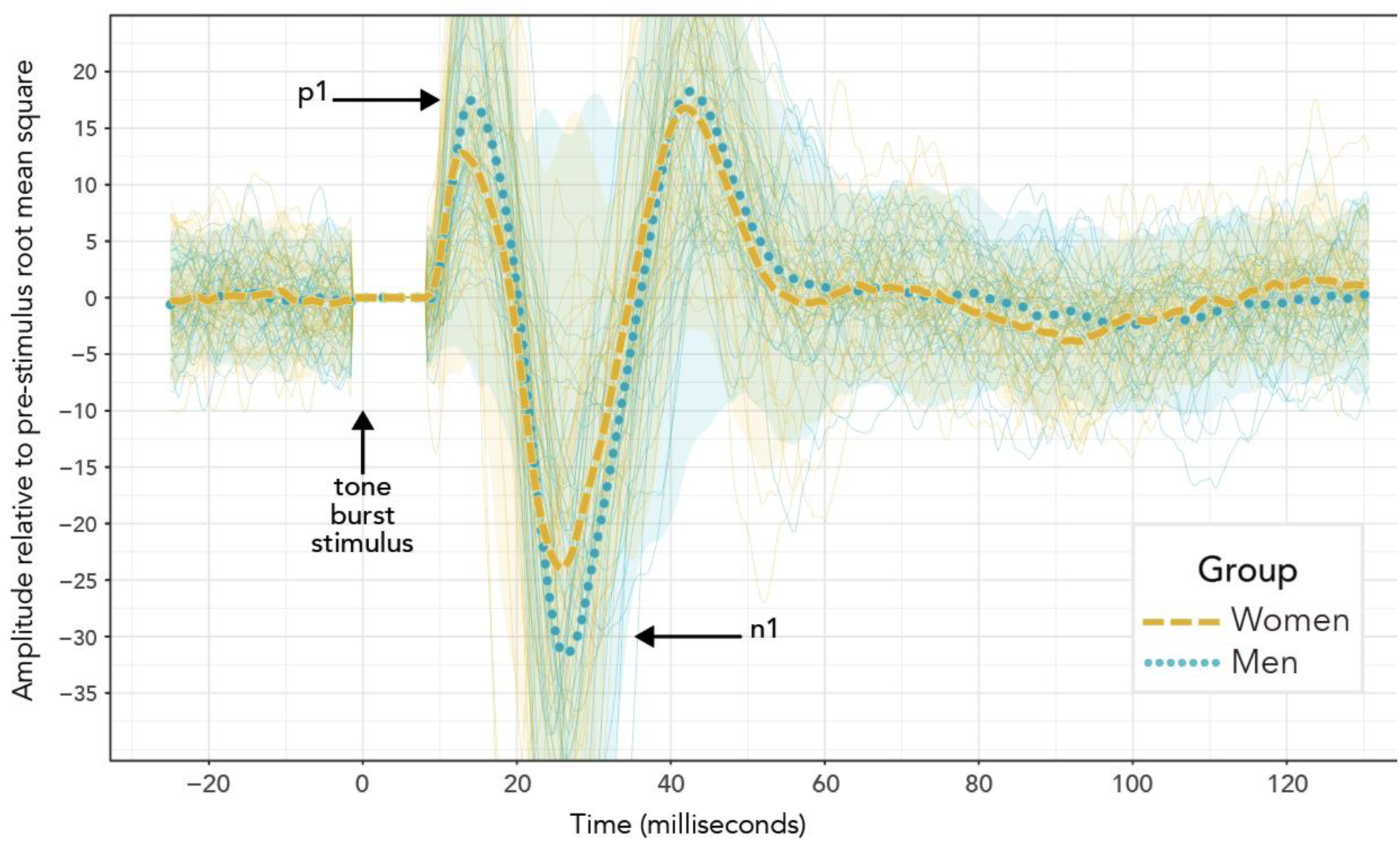
VEMP grand averages recorded following 500 Hz tone bursts presented using bone conduction at 40 dB HL. Visual inspection of the waveforms suggests a difference in p1-n1 amplitude, but not p1-n1 latency. However, these data do not enable a valid statistical comparison, since there were uneven presentation counts across women and men. Also shown are individual sequences for each group (more lightly drawn traces than the grand averages) along with 95% confidence intervals (shaded areas). A Welch’s unequal variances t-test was conducted on peaks and troughs calculated per sequence, once repeated measures at 40 dB HL had been averaged across each participant. This showed no group difference in VEMP p1-n1 amplitude (p= 0.53; mean (women) = 48.9; mean (men) = 55.0; 95% CI -25.7, 13.4; t = -0.6; df = 45) and a statistically significant group difference in which VEMP p1-n1 latency was 2.4 ms shorter in women than men (p = 0.01; mean (women) = 10.3; mean (men) = 12.7; 95% CI -4.3, -0.63; t = -2.7; df = 42). The actual statistical comparison between women and men used linear mixed-effects regression modelling, with a much wider range of stimulus presentation levels. Statistical analyses are reported in more detail in section 3, and discussed in section 4.

For other stimulus levels, peaks and troughs were identified using a similar process to that just described for the initial 40 dB HL sequence. A difference was that the trough from the initial 40 dB HL sequence was used as an anchor for trough detection for remaining sequences on a per participant basis. A null identification of peaks and trough occurred (no result was returned from the script) if the p1-n1 amplitude was less than 1.65 times the pre-stimulus RMS for the sequence of 300 repetitions being evaluated.

The script was verified through visual inspection of waveforms for the entire data set collected. Creation of the procedure was via an iterative process, with adjustments made to some of the parameters which have been described prior to re-running the script. Visual inspection showed the final script identifying peaks and troughs accurately. Identification made by the script was final – data points were not removed or adjusted manually.

Data were converted into a response level (RL) scale by taking the logarithm of p1-n1 amplitude:

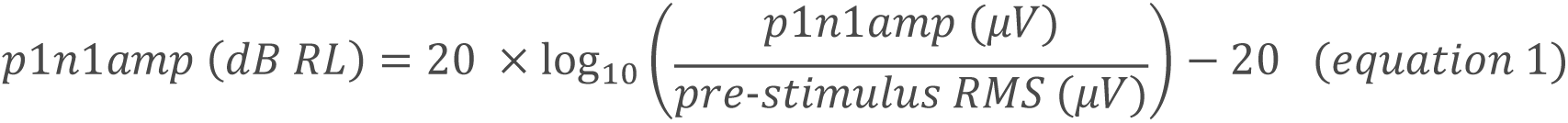

Zero dB RL corresponds to a projected VEMP threshold (this projection is unlike the VEMP thresholds identified in clinical procedure). The transformation is similar to that for the dB SPL scale popular for sound pressure levels, and its frequency-adjusted HL variant, in which 10 dB increase corresponds to an approximate doubling perceptually.

### 2.5 Confounders in VEMP measurement

Various precautions were taken to minimise potential confounders in VEMP measurement. These mainly addressed measurement of VEMP p1-n1 amplitude, but will also have increased accuracy in measuring VEMP p1-n1 latency.

#### 2.5.1 Stimulus level

VEMP p1-n1 amplitude has been found to increase with stimulus level (Colebatch et al., 2016). Linear mixed-effects regression modelling was conducted based on this relationship, with between group comparisons calculated from VEMP growth rate.

#### 2.5.2 Neck tension

To record a cervical VEMP, the sternocleidomastoid muscle (SCM) must have a tension greater than that at resting state (Colebatch, Rosengren & Welgampola, 2016). However, VEMP p1-n1 amplitude has been found to increase with SCM tension (Ochi et al., 2001). To prevent SCM tension acting as a confounder, its variation was limited. This was achieved in several ways. One was by instructing participants to maintain a constant biofeedback target by pushing their foreheads against a padded head bar (figure 2). Another was by measuring pre-stimulus SCM tension so that it could, if necessary, be included as a covariate. A third was to assess duration of testing, in case fatigue contributed to variation in SCM tension.

#### 2.5.3 Age

Age matching between participants was used to control for a decrease in VEMP p1-n1 amplitude with age (Nguyen et al., 2010; Colebatch et al., 2013) and a prolongation of VEMP p1 and n1 latencies with age (Brantberg et al., 2007; Macambira et al., 2017).

#### 2.5.4 Crossed response

Cervical VEMPs have been found to be predominantly ipsilateral, however components have sometimes been observed contralaterally (Colebatch & Rothwell, 2004; Ashford et al., 2016). Delivery of binaural stimuli with a bone conductor limited variation due to any between participant difference in the contralateral nature of VEMPs, because both ipsilateral and contralateral components of the VEMP from each ear were present at both SCM muscles. The arrangement was not perfect. The bone conductor placement at the mastoid introduced an asymmetry, with an attenuation of approximately 3–5 dB intracranially for the 500 Hz tone burst used (Stenfelt, 2012). However, the asymmetry was consistent per participant.

#### 2.5.5 Sternocleidomastoid physiology

VEMP amplitude may be influenced by the size of the SCM and subcutaneous fat (Chang et al., 2007; Bartuzi et al., 2010). This study did not appraise volume of SCM and subcutaneous fat, however any variations will have been minimised by the normalisation procedure, pairing on age and sex, and use of amplitude growth parameters in the between group comparisons.

The exact positioning of electrodes along the SCM (e.g. positioning as a function of proximity to the belly of the SCM) has been found to correspond to substantial variation in VEMP measurements (Rosengren et al., 2016; Ashford et al., 2016). Studies in which VEMP latency increased with neck length (Chang et al., 2007; Wang et al., 2008) have underscored the importance of electrode placement, due to the increased scope for inaccuracy while placing an electrode over the belly of the SCM on a longer neck. Placing electrodes a small distance away from the belly of the SCM has been found to prolong VEMP latency measurements (Rosengren et al., 2016).

Placement of electrodes followed a clinical palpation technique (BSA, 2012) aiming for a consistent position above the SCM belly. However, it is difficult to escape the conclusion that with a clinical VEMP technique, “small errors in electrode placement with respect to the motor point are likely to be inevitable” (Rosengren et al., 2016). The palpation technique was consistent across female and male participants meaning that, based on the central limit theorem, inaccuracy in electrode placement will have decreased as a function of participant count. Participant count will be discussed further in section 4.1. Electrode placement will be one of the factors contributing to the disturbance in the statistical model to be described in figure 7, as will variation in neck length. The study of Brantberg et al. (2007), whose count of 1000 participants was an order of magnitude greater than all but one of the studies in table 1, should be preferred to other studies when electrode positioning is considered.

#### 2.5.6 Blood flow

Blood has electromagnetic properties (Beving et al., 1994; Abdalla, 2011). It follows that electromagnetic field variation due to blood flow could add noise during electromyographic recording. The active electrode placement for cervical VEMPs was directly above the carotid artery, such that effect of blood flow on individual epochs could have been appreciable. However, the effect of blood flow was mitigated firstly by the large size of the VEMP response, and secondly by the averaging process applied to the electromyographic recordings. The stimulus delivery rate was 5.1 per second, whilst resting state pulse rates are approximately one per second. Over the approximately one minute recording time for an averaged sequence of 300 epochs, variations in the electromyographic recording due to carotid artery blood flow will have largely cancelled out, such that noise due to blood flow will not have appreciably affected within or between participant comparisons.

## 3 Results

### 3.1 VEMP p1-n1 amplitude

The histogram in figure 4 shows counts of VEMP p1-n1 amplitude measurements sorted into female and male participants. The histogram does not show detail of participant or stimulus level. Inclusion of repeated measurements in the histogram means that it is not appropriate for statistical comparisons. However, the presentation count was approximately equal per participant, and was carried out over approximately the same stimulus range. Therefore the histogram gives an indication of distribution for each group. Both the female and male groups appear to have approximately normal distributions. It is apparent that slightly more data have been collected from men than from women. There is no suggestion of difference between the means of the distributions.

**Figure 4:**
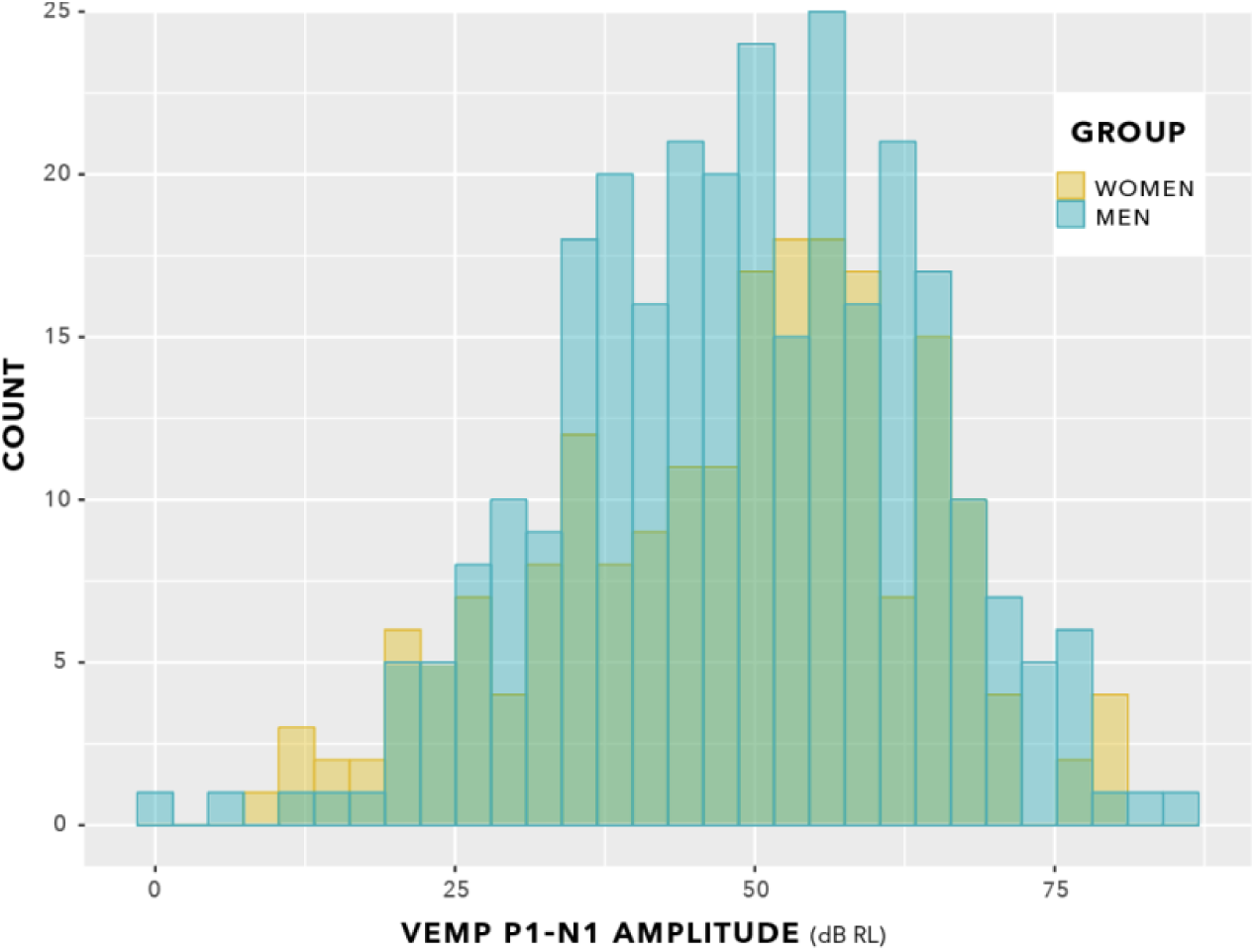
VEMP p1-n1 amplitude histogram. Indication is of approximately normal distributions for the female and male groups. There is no suggestion of a difference in means of the distributions. Slightly more data were collected from men than from women.

The box plot in figure 5 provides an alternative view of the data in figure 4. Again, there is no suggestion of group difference, however this was verified by statistical analysis.

**Figure 5:**
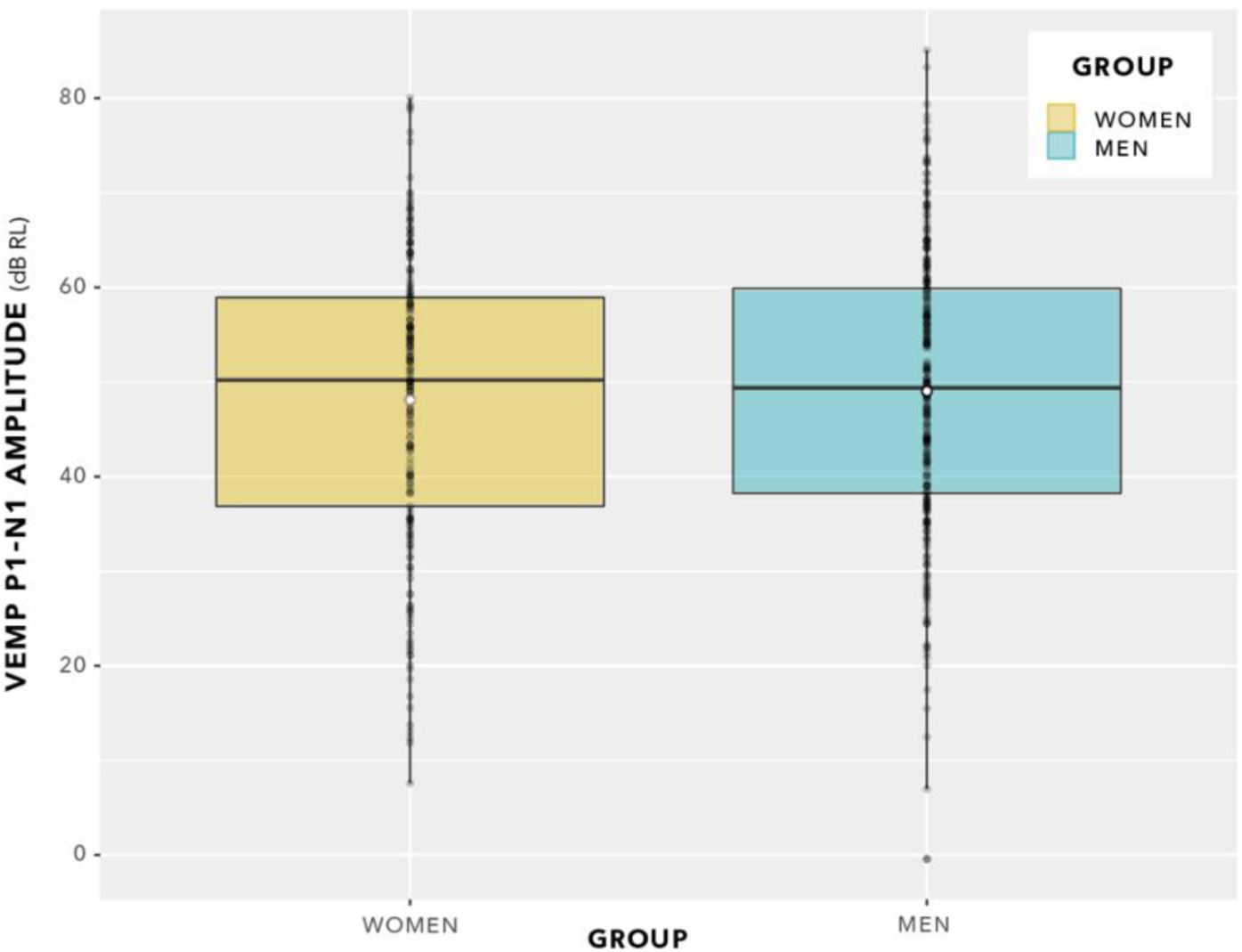
VEMP p1-n1 amplitudes. Since there are repeated measures per participant, this is not a valid statistical comparison. However, there is no indication of a statistically significant group difference. Statistical analysis was via linear mixed-effects regression modelling, and is described in the main text.

The distributions in figure 5 do not enable a valid statistical comparison, because they include repeated measurements. Statistical comparison was achieved through linear mixed-effects regression modelling, using stimulus level as a predictor, as illustrated in figure 6. This is a log-log graph, since both the abscissa (stimulus level) and the ordinate (VEMP p1-n1 amplitude) are transformed into logarithmic scales. Thus, a power relationship is apparent between stimulus level and VEMP p1-n1 amplitude. As in figures 4 and 5, figure 6 shows no suggestion of group difference.

**Figure 6:**
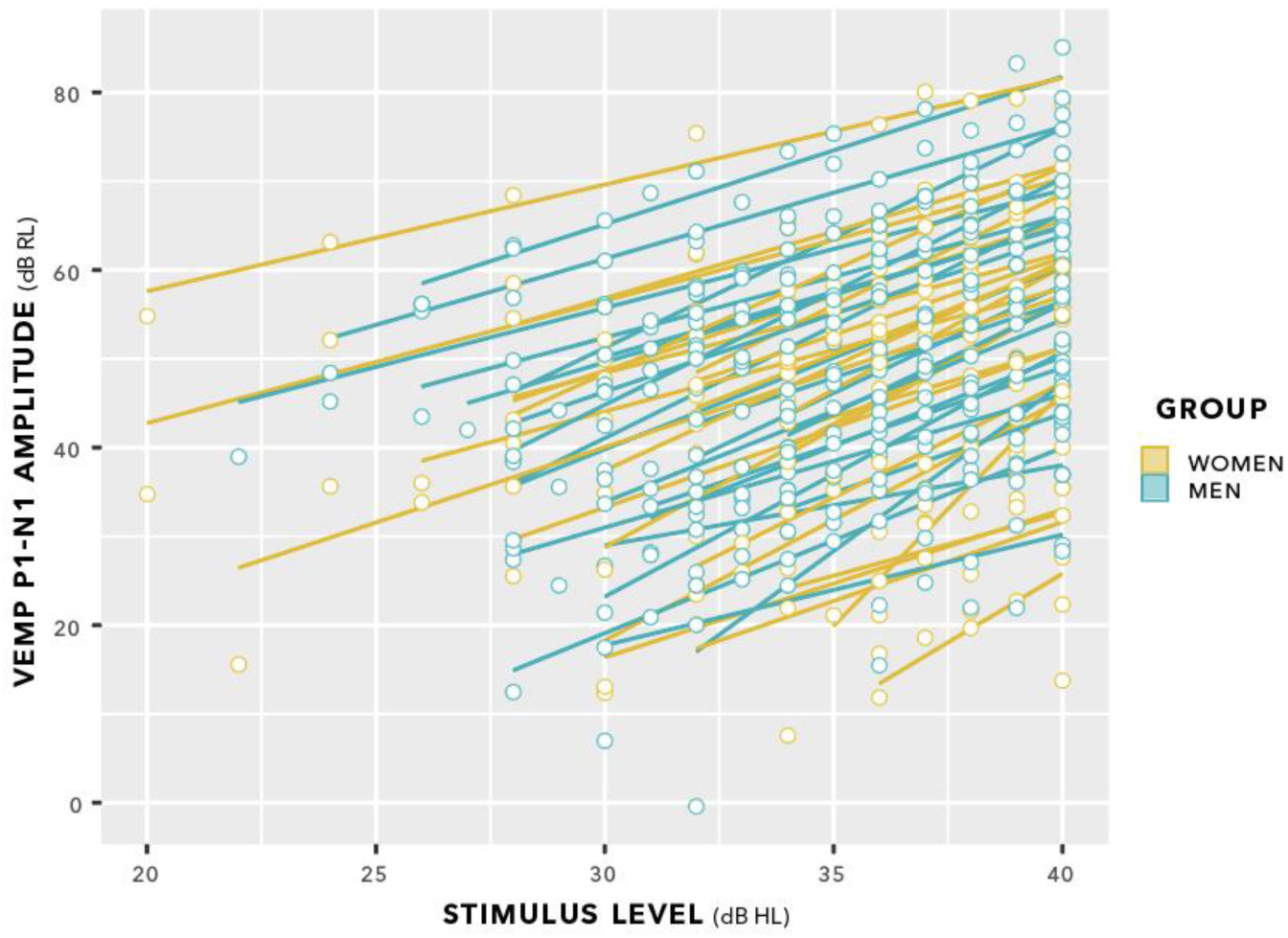
VEMP p1-n1 amplitude versus stimulus level. Individual lines of best fit are per participant.

The initial statistical model for VEMP p1-n1 amplitude is shown in figure 7. It included potential confounders, as per the review in section 2.5.

**Figure 7:**
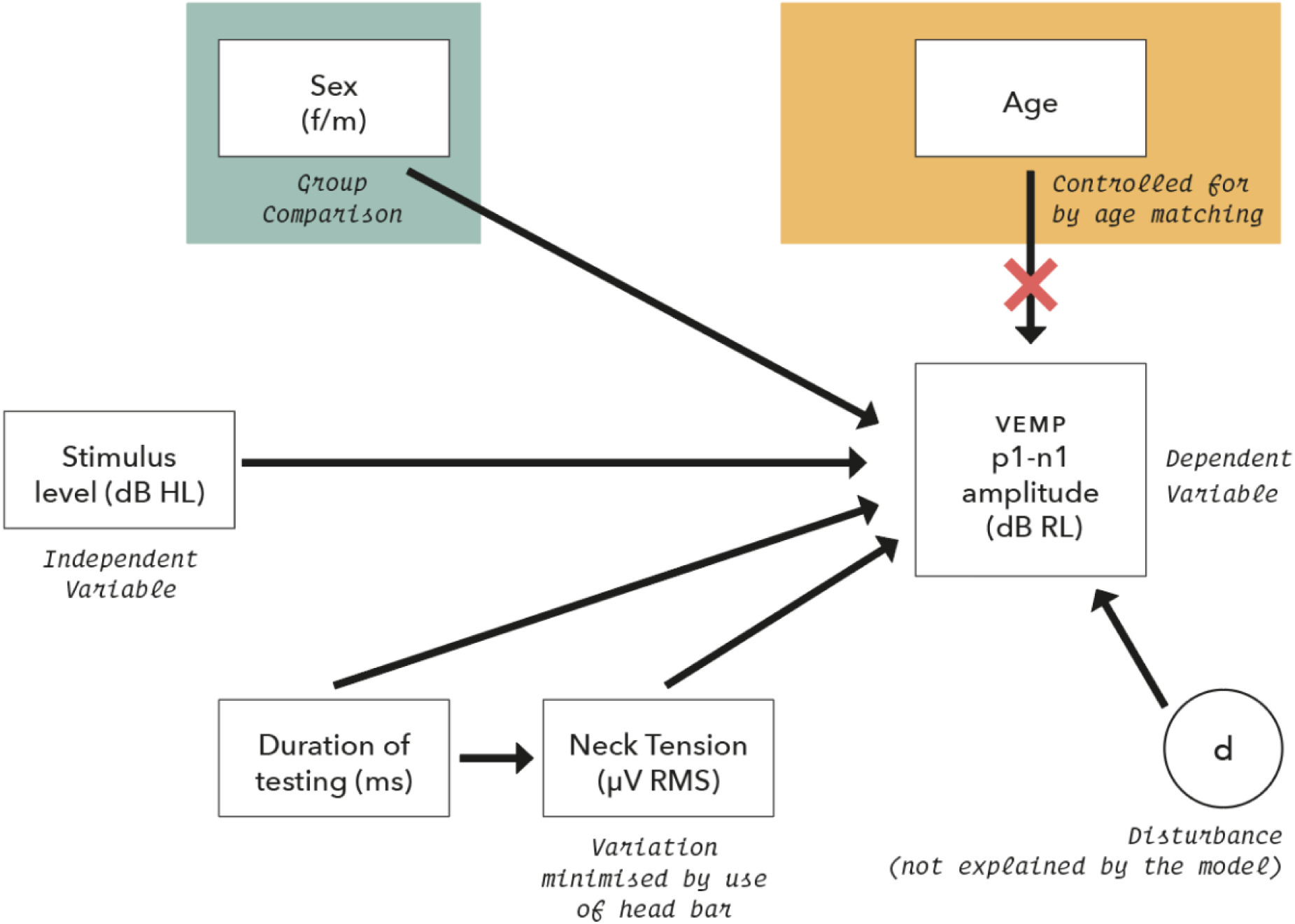
Initial model for statistical analysis of VEMP p1-n1 amplitude measurements. Potential confounders were reviewed in section 2.5, and were assessed as part of the model where possible.

Linear mixed-effects regression modelling has been shown to entail a trade-off. There is an increased possibility of type I error when data from all participants are assigned the same slope but can have varying intercepts, versus lower statistical power when both slope and intercept can vary per participant (Barr et al., 2013; Matuschek et al., 2017). The slopes in figure 6 have a mean value of 1.89 with SD 0.25, so a fixed slope appeared a reasonable assumption. Use of a fixed slope model was further supported by the precursor to this study (Gattie et al., 2021), which assessed fixed and varying slope linear mixed-effects regression models (Winter, 2019) for VEMP p1-n1 amplitudes with participants who did and did not stutter, and found that a fixed slopes model was most appropriate. Accordingly, a fixed slope model was used in analysis of the current study, with a form as follows:

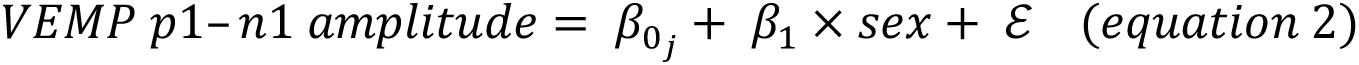

VEMP p1-n1 amplitude was conditioned on whether participants were female or male, with ß_0_ as intercept (varies with participant, j) and ß_1_ as a fixed slope of increase in VEMP p1-n1 amplitude with stimulus level. Statistical analysis was conducted with the lme4 package (Bates & Maechler, 2020) in R (Team, 2020). Effect size (Cohen’s d) was calculated from mixed model t statistics with the EMAtools package for R, version 0.1.3 (Team, 2020). Conditional R^2^ was calculated according to Nakagawa et al. (Nakagawa et al., 2017) using the MuMIn package, version 1.43.17 (Team, 2020).

Likelihood ratio comparisons to a nested model in which sex of participant as a predictor was absent was used to calculate p values. There was no significant difference between models evaluated (Chi-Squared (1) = 0.41, p = 0.52), indicating that VEMP p1-n1 amplitude did not depend on the sex of the participant. Conditional R^2^ was evaluated as 0.92. Overall results are shown in figure 8.

**Figure 8:**
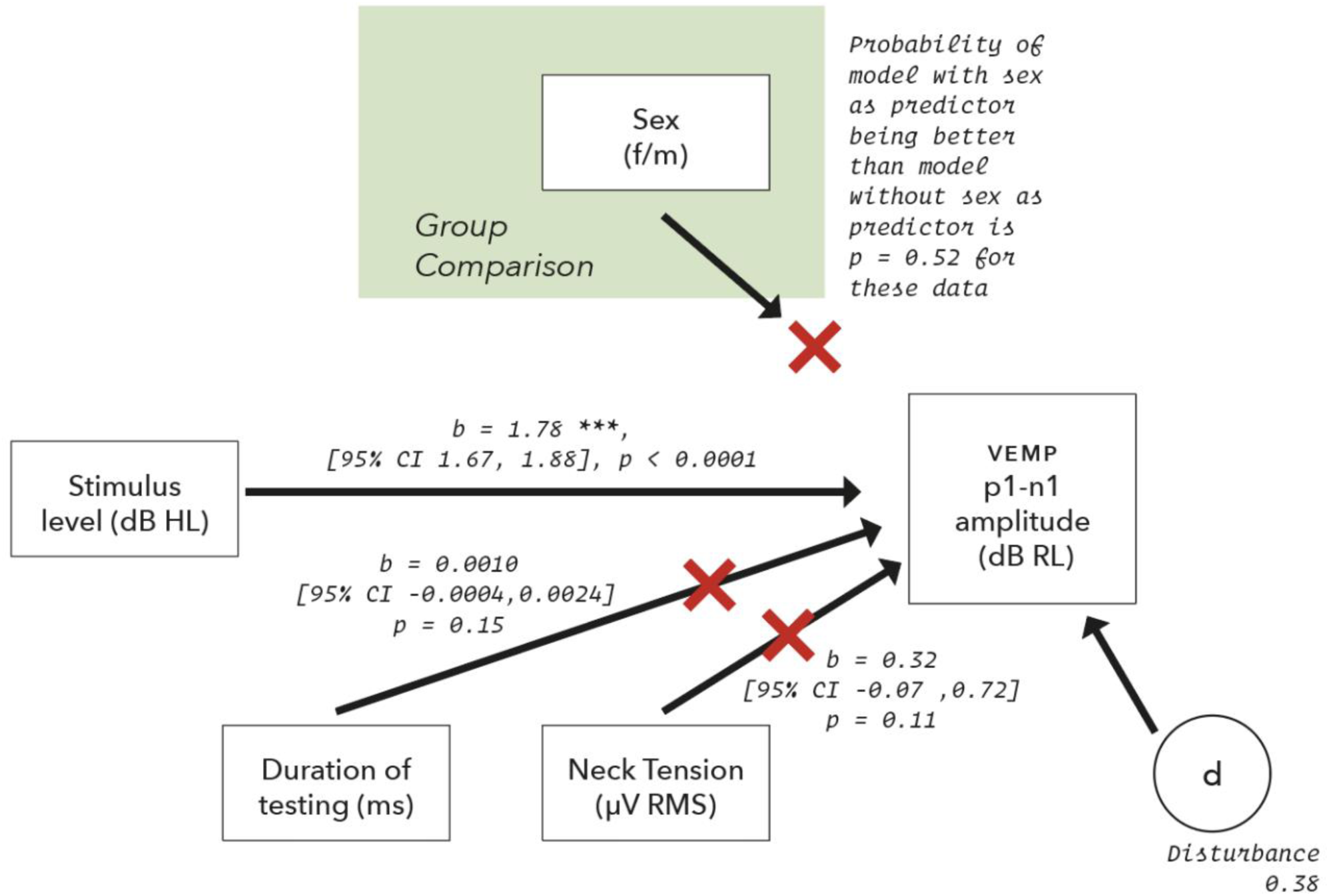
The model for VEMP p1-n1 amplitude described in figure 7, updated following data collection. Only stimulus level predicted VEMP p1-n1 amplitude.

Both time elapsed and pre-stimulus RMS had *p* > 0.1, and a small effect on p1-n1 amplitude relative to stimulus level. They were therefore removed from the final model, which is shown in figure 9. Only stimulus level predicted VEMP p1-n1 amplitude. Data used in this analysis are available in the supplementary material.

**Figure 9:**
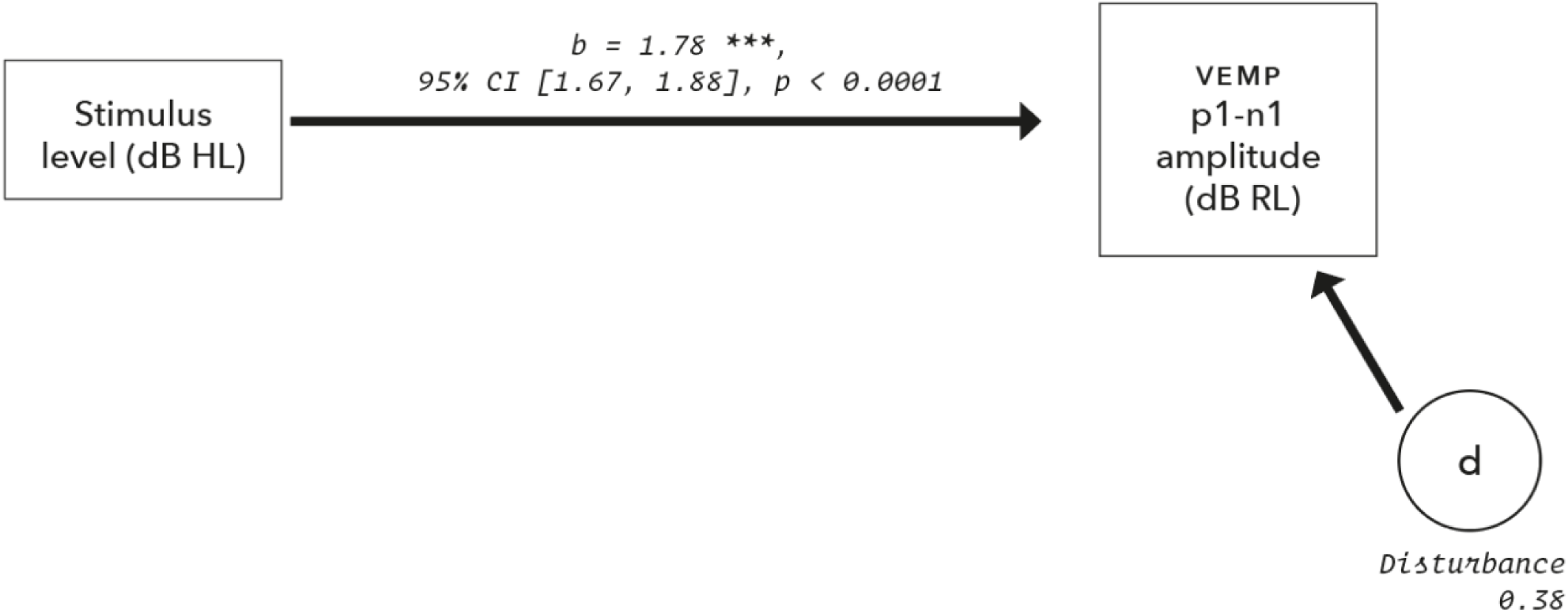
Final model for VEMP p1-n1 amplitude.

### 3.2 VEMP p1-n1 latency

Figure 10 shows latencies collected across all participants and all stimulus levels, including repeat measurements. Data appear normally distributed, with suggestion of a group difference.

**Figure 10:**
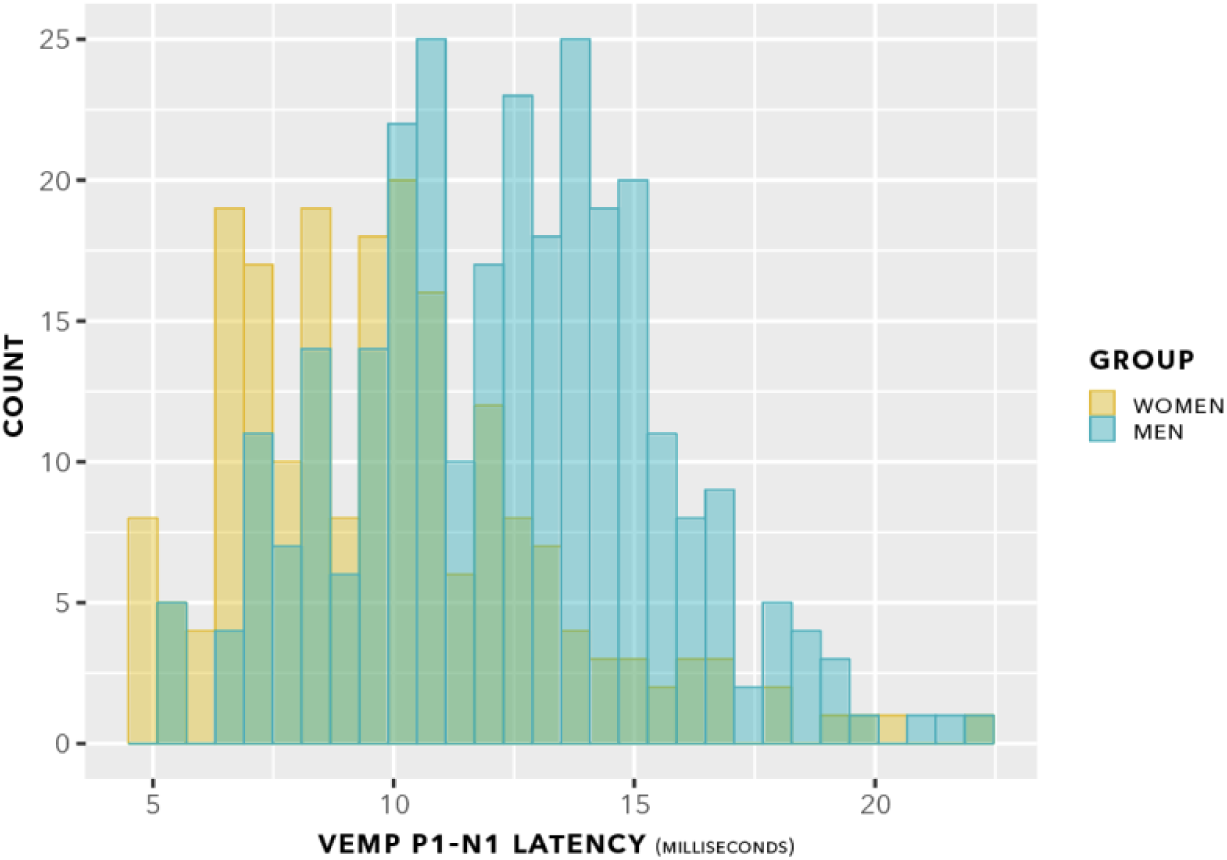
VEMP p1-n1 latency histogram. Indication is of approximately normal distributions for the female and male groups, with some suggestion of a difference in means of the distributions.

The data from figure 10 are shown in a box plot in figure 11. There appears to be a group difference.

**Figure 11:**
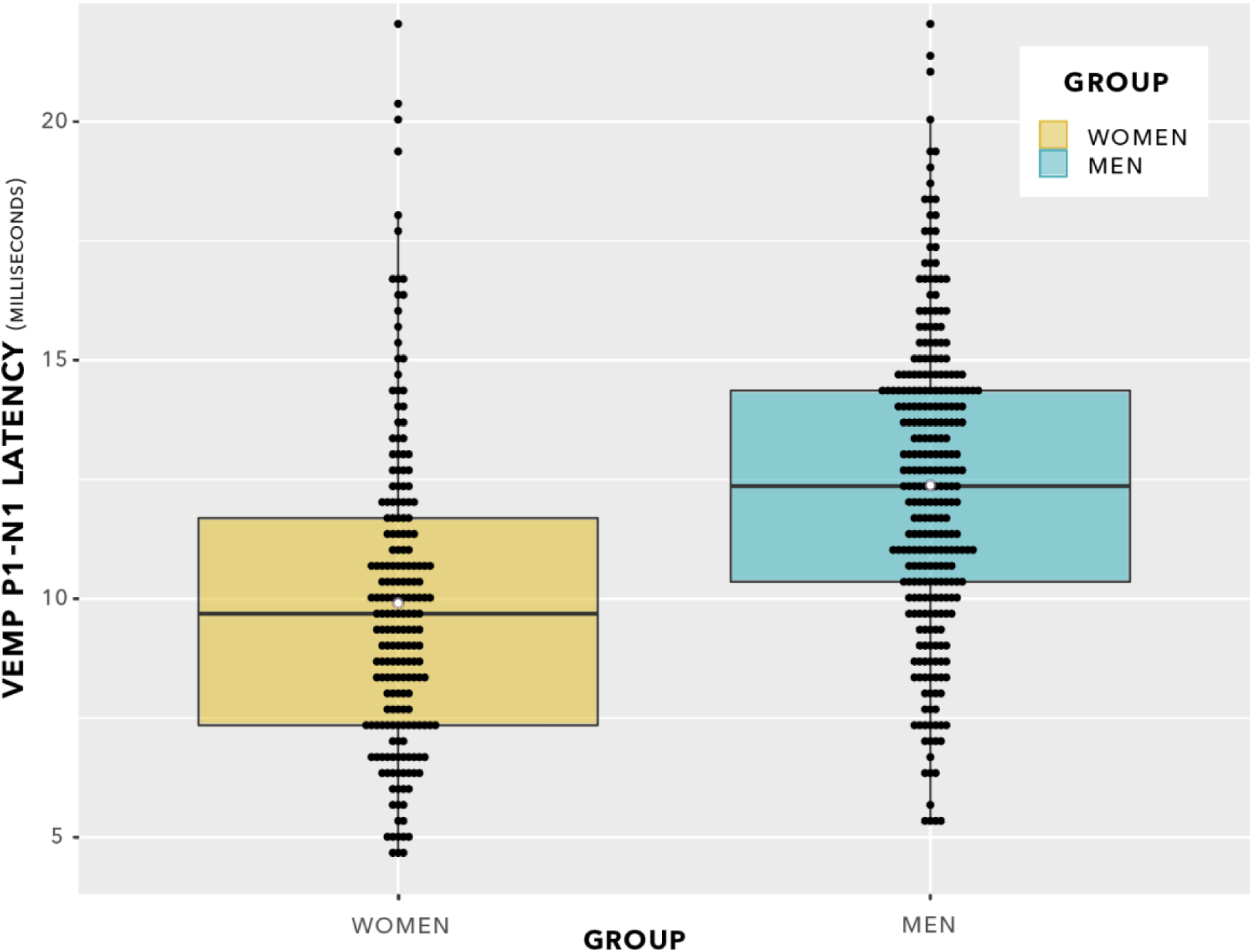
VEMP p1-n1 latencies. The even spacing on the ordinate was due to the 3 kHz sampling rate. VEMP p1-n1 latencies are grouped in steps of 0.33 ms, which was the maximum temporal resolution of the Eclipse when using VEMP protocols. To indicate when several data points were recorded at the same VEMP p1-n1 latency, data points have been plotted using the beeswarm feature in ggplot (R Core Team, 2020). This feature has introduced variation along the abscissa corresponding to the quantity of data present. Mean values for each group are shown as white data points. Although the box plot suggests there will be a group difference in VEMP p1-n1 latency, it contains repeated measurements per participant. As such, the box plot does not entirely reflect data appropriate for use in statistical analysis. However, the repeated measurements per participant could be averaged, enabling statistical comparison using a Welch’s unequal variances t-test. Such a t-test was conducted, with the finding that p1-n1 latency was 2.4 ms shorter in women than in men. The finding was statistically significant at p = 0.0026 (95% CI -0.89, -3.96; t stat -3.18, df 46, mean (women) 10.0, variance 7.5; mean (women) 10.0; mean (men) 12.4; variance 6.4). This compares with p = 0.0020 for the linear mixed-effects modelling comparison, as described in the main text.

Statistical evaluation was carried out by linear mixed-effects modelling. First, the effect of stimulus level on VEMP p1-n1 latency was appraised. The following models were compared:

model_null: latency ∼ 1 + group + (1| participant)
model_diff: latency ∼ stimulus + group + (1| participant)

This comparison showed no significant difference between models (chi squared (1) 0.06, p = 0.80). The indication was of no effect of stimulus level on p1-n1 latency. This is illustrated in figure 12. Further comparisons indicated no effect of duration of testing or of neck tension.

**Figure 12:**
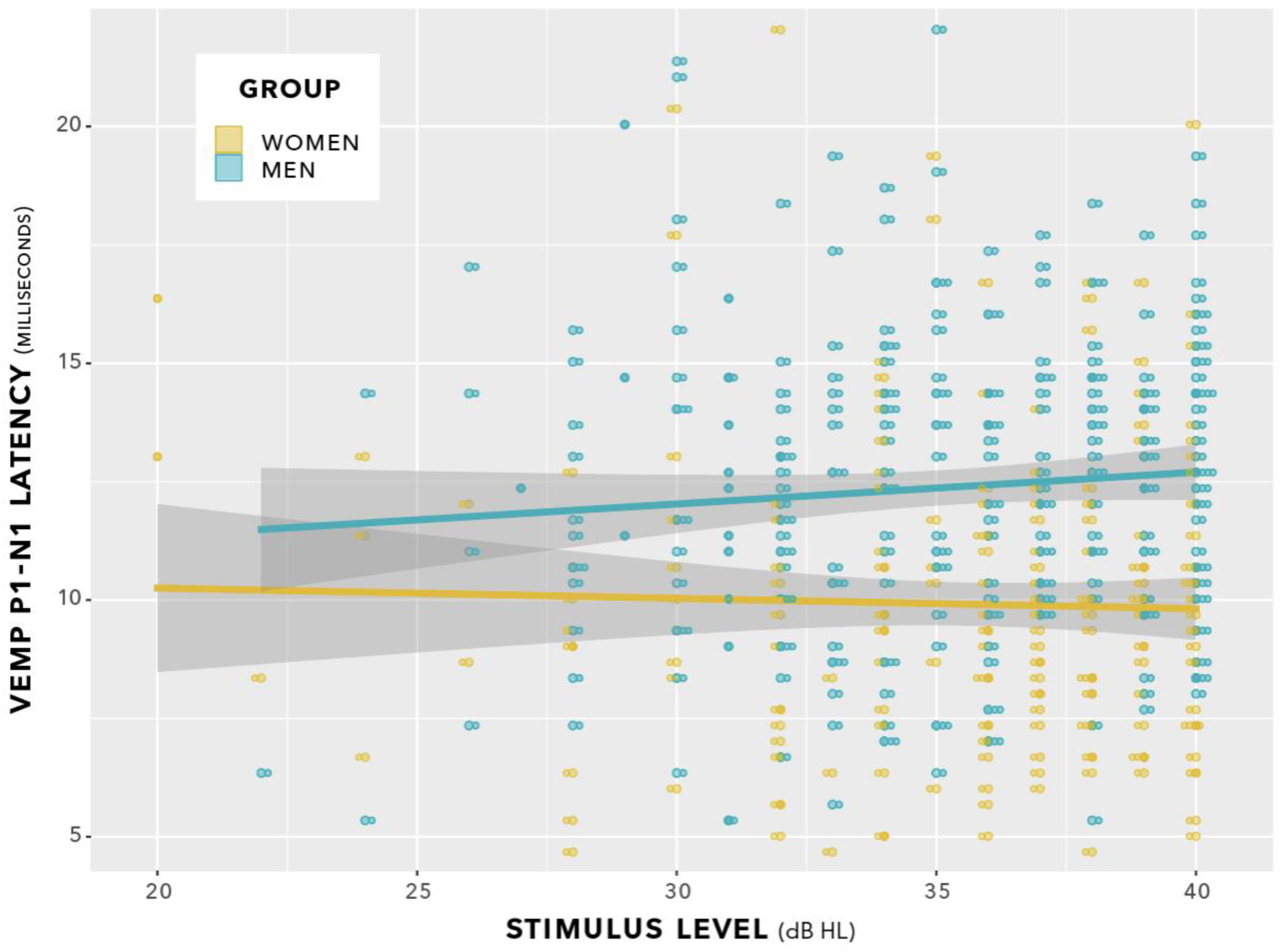
VEMP p1-n1 latencies versus stimulus level for women and men. There was no interaction. Sampling resolution of the Eclipse meant that data points were grouped in steps of 0.33 ms (see note at figure 11). To provide an indication of when data from both women and men were recorded at the same stimulus level and VEMP p1-n1 latency, data points have been plotted using the beeswarm feature in ggplot (Team, 2020). This feature introduced variation along the abscissa corresponding to the quantity of data present. The variation does not reflect stimulus level used. Stimulus level was always an integer multiple of 1 dB.

Next, p-values were generated by likelihood ratio comparisons between the following models.

model_null: latency ∼ 1 + (1| participant)
model_diff: latency ∼ 1 + group + (1| participant)

There was a significant difference between models (chi squared (1) 9.6, p = 0.002).

The data showed that VEMP p1-n1 latency was shorter for women than for men by 2.43 ms (95% CI [–0.93, –3.92], chi squared (1) 9.6, p = 0.0020). Data used in this analysis are available in the supplementary material.

### 3.3 Absolute p1 and n1 VEMP latencies

Absolute p1 and n1 VEMP latencies were evaluated, using linear mixed-effects modelling similar to that already described for p1-n1 interpeak latency. These analyses are made available for readers wishing to compare results from this study with the prior studies (see table 1) which have reported absolute p1 and n1 VEMP latencies. These additional analyses are presented for exploratory purposes only. Analyses of absolute VEMP p1 and n1 latencies were not pre-registered, multiple corrections have not been made enabling such additional analyses, and results from the additional analyses are not a focus of discussion in this report.

For women, the absolute p1 latency was 15.3 ms [95% CI 14.5, 16.1] and the absolute n1 latency was 25.3 ms [95% CI 24.3, 26.3]. For men, the absolute p1 latency was 14.6 ms [95% CI 14.2, 15.0] and the absolute n1 latency was 27.0 ms [95% CI 26.1, 27.8]. There was no group difference in p1 latency having statistical significance at the p ≤ 0.05 level, although a statistically insignificant and uncorrected p1 prolongation of 0.74 ms in women compared to men was observed (p = 0.11, chi squared (1) = 2.54, standard error = 0.46). For n1 latency, there was a statistically significant group difference at the p ≤ 0.05 level. The n1 trough occurred 1.69 ms earlier in women than in men (p = 0.015, chi squared (1) = 5.90, standard error = 0.67). Even with a Bonferonni correction, this group difference would remain statistically significant at p ≤ 0.05.

## 4 Discussion

### 4.1 Comparison to prior studies

This study found that VEMP p1-n1 latency was shorter in women than men by 2.4 ms (95% CI [– 0.9, –3.9], chi squared (1) 9.6, p = 0.0020). The VEMP p1-n1 latency differences equates to 21% of the mean 11.4 ms VEMP p1-n1 latency, which was measured across women and men. The study also found that there was no sex difference in VEMP p1-n1 amplitude.

Prior studies of sex difference in p1-n1 latency and amplitude were summarised in table 1. Of the 14 studies other than the current one, 10 found no sex difference in p1-n1 latency. The remaining four studies consistently found women with shorter p1 or n1 latencies than men. However, when the p1-n1 latency difference is considered, the picture is as mixed as for p1-n1 amplitude. Two studies found a shorter p1-n1 latency in women than in men (Lee et al., 2008; Brantberg et al., 2007) and two studies found a longer p1-n1 latency in women than in men (Brantberg & Fransson, 2001; Li et al., 2015).

For VEMP amplitudes, 10 of 14 studies found no sex difference. Of the remainder, two found larger p1-n1 amplitudes in women (Welgampola & Colebatch, 2001; Lee et al., 2008), one found larger p1-n1 amplitudes in men (Shahnaz & David, 2021), and one found larger p1-n1 amplitudes in men but in the right ear only (Silva et al., 2016). For VEMP p1-n1 amplitudes, it is not just that a minority of studies found a sex difference, but moreover that when a sex difference was found, the direction of the difference was inconsistent.

There are at least two interpretations of the prior research. In the first, VEMP has no genuine difference in p1-n1 latency or p1-n1 amplitude between women and men. The rationale for this explanation is that in half of the studies in Table 1 no sex difference in VEMP measures was found, and in the other half there was no consistent direction of sex difference in either p1-n1 latency or p1-n1 amplitude. The occasional differences established might, for example, have been due to sampling variation, confounders specific to individual studies, or a combination of both. An alternative explanation is that prior studies were underpowered. This second explanation is compatible with aspects of the first explanation (e.g. occasional findings of sex difference due to sampling variation or study-specific confounders), but it adds the detail that a genuine sex difference in VEMP measures could have been missed due to a lack of sensitivity in the tests applied. Three aspects of the current study support this alternative explanation.

Firstly, use of the head bar and biofeedback in the current study provided greater control of SCM tension than the head turn or head raise procedures in prior studies. This has greater importance for measurement of p1-n1 amplitude than for p1-n1 latency, since p1-n1 amplitude is proportional to SCM tension (Ochi et al., 2001). Physiological effects related to SCM tension are discussed in detail in section 4.2. The indication from the current study is that there is no sex difference in VEMP p1-n1 amplitude.

Secondly, age matching between sexes was more exact in the current study than in prior studies. Participants in the current study were aged between 18–21 years, plus one 16 year old, with a mean age of 19.5 and SD of 0.7. Age matching between sexes in prior studies was far less exact and is sometimes not even reported. Less exact age matching introduced a confounder, since across the lifespan p1-n1 amplitudes are attenuated by a factor of 2 (Welgampola & Colebatch, 2001; Brantberg et al., 2007) and there is variation of between 1 and 3 ms in p1 and n1 latencies (Brantberg et al., 2007; Macambira et al., 2017). Thus, the failure to establish a sex difference in some prior studies may be because a genuine sex difference was obscured by an age effect which was not adequately controlled for. Equally, when a sex difference was established, sometimes with a different direction of fit to the current study, or in terms of amplitude rather than latency, the finding may in fact have reflected a variation in age rather than sex between the groups which were compared.

Thirdly, the use of BC stimuli in the current study enabled collection of a far greater amount of data per participant than any of the studies described in table 1. Typically the number of stimulus presentations was an order of magnitude greater – a mean of 3044 presentations per participant in the current study, versus between 200 and 300 per participant in prior studies. The effect of collecting a greater number of samples of VEMP responses per participant will have been to decrease sampling variation in the within participant measure. Reduction of sampling variation in the within participant measure in turn reduces the chance of reporting an apparently statistically significant, but in actuality spurious, difference in the between group comparison, and increases the chance of reporting a genuine group difference.

The studies described in table 1 were not primarily designed for appraisal of sex difference in VEMP measures and, from the considerations just described, may have been underpowered to do so. This possibility could be appraised with a power analysis based on data from the current study. There are serious caveats when interpreting post hoc power analyses (Hoenig & Heisey, 2001; Lakens & Evers, 2014). These are compounded in the current study by the difficulty of estimating power in linear mixed-effects regression modelling (Kumle et al., 2021). A standard solution to the latter difficulty is to implement Monte Carlo simulation, however even with stock routines (Green & MacLeod, 2016) the programming overhead would have been beyond scope for this discussion section. Fortunately, there was a more straightforward solution. As described in the caption for figure 11, averaging VEMP p1-n1 latency measurements per participant then running a Welch’s unequal variances t-test gives near-identical results to the linear mixed-effects regression analysis. Use of t-tests enables adoption of the power analysis routines described by Cohen (1969). Such analyses were carried out in table 2, with power calculations according to G*Power (Faul et al., 2007). The effects of fewer stimulus presentations, and fewer participants, were both modelled. Results are presented graphically in the form of 95% confidence intervals in figure 13. Data used in these analyses are available in the supplementary material.

**Figure 13:**
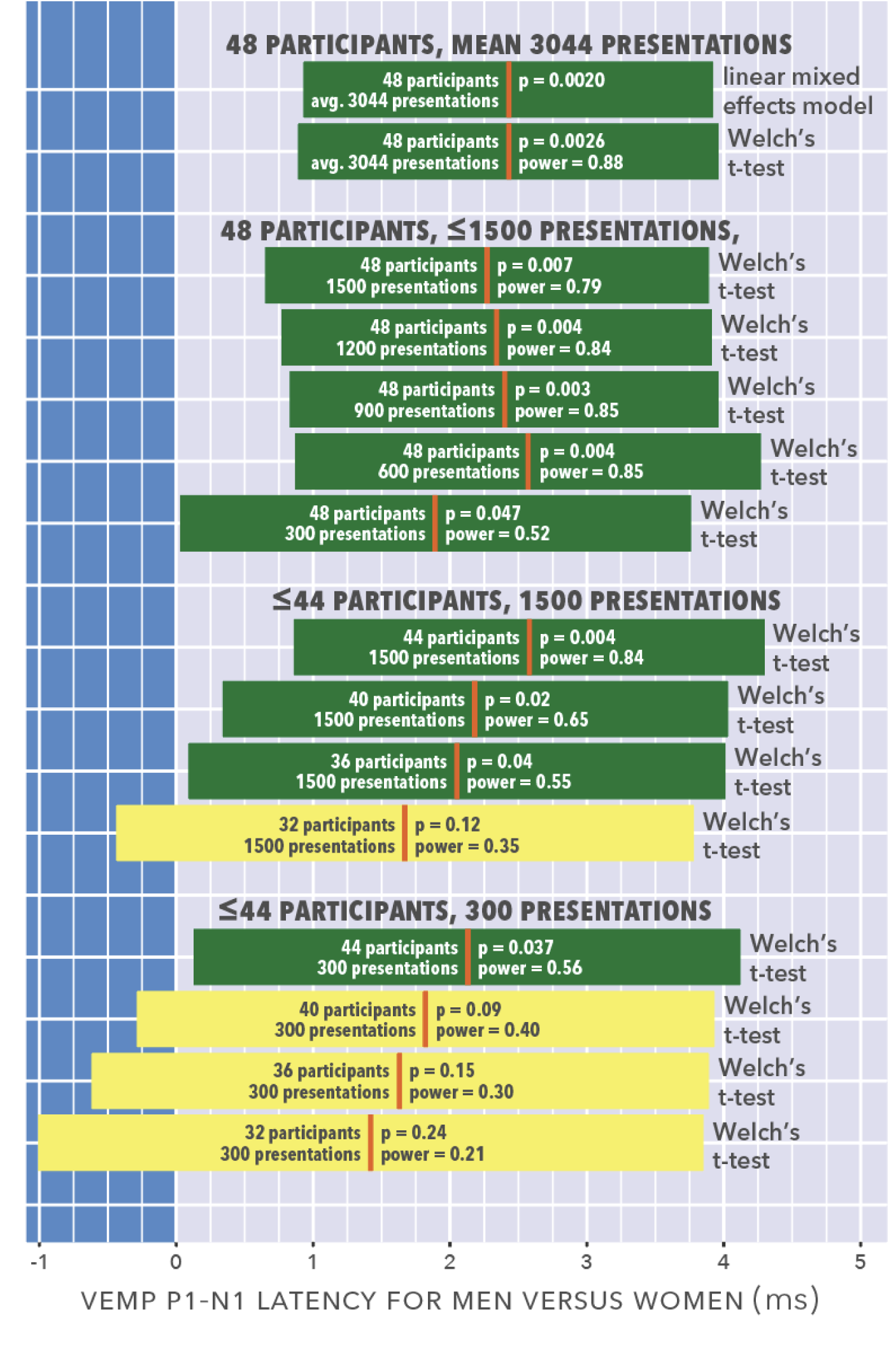
95% confidence intervals for the analyses shown in table 2. Participant counts are balanced (e.g. 48 participants are 24 women, 24 men). Group differences are shown as prolongation in men’s VEMP p1-n1 latencies compared to those of women. The mean latency difference is shown for each condition as a vertical orange line, with horizontal bars either side showing the 95% confidence interval. When the 95% confidence interval crosses zero, the data analyses in the simulations did not establish a statistically significant group difference. Such analyses are depicted in a lighter colour (yellow), whereas analyses for which a statistically significant group difference could be established are depicted in a darker colour (green).

**Table 2:**
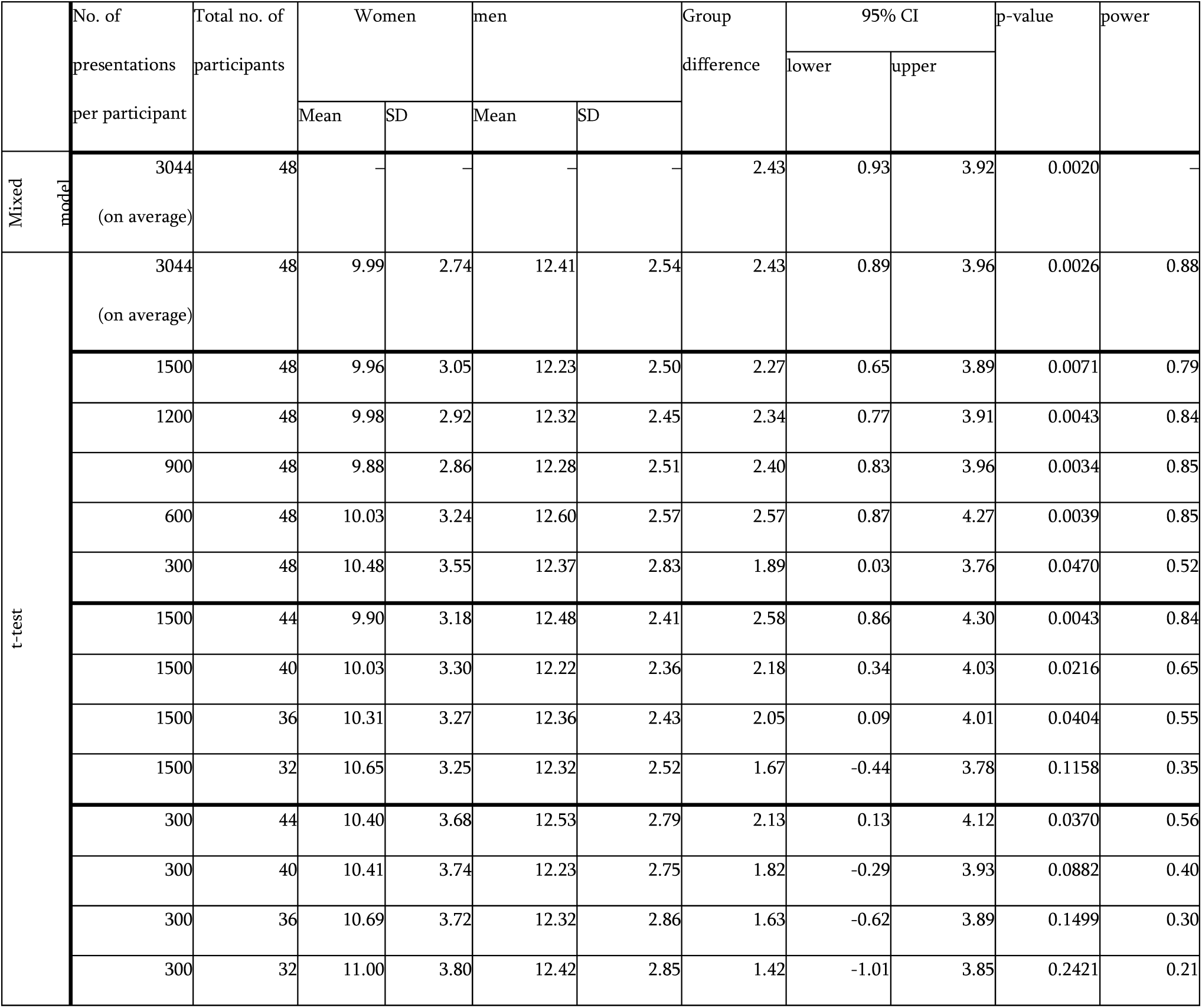
Modelling of the effect of presenting fewer stimuli, and/or testing fewer participants. The results presented in section 3.2 used linear mixed-effects modelling, with an average of 3044 presentations for each of 48 participants (24 women, 24 men). As described in the caption to figure 11, an analysis with near-identical results is possible using a Welch’s unequal variances t-test. For the simulation, t-tests were recalculated after reducing either participant count or number of presentations. In all simulations, the data which were collected last were removed prior to reanalysis.

|  | No. of presentations per participant | Total no. of participants | Women |  | men |  | Group difference | 95% CI |  | p-value | power |
| --- | --- | --- | --- | --- | --- | --- | --- | --- | --- | --- | --- |
|  |  |  | Mean | SD | Mean | SD |  | lower | upper |  |  |
| Mixed model | 3044<br>(on average) | 48 | — | — | — | — | 2.43 | 0.93 | 3.92 | 0.0020 | — |
| t-test | 3044<br>(on average) | 48 | 9.99 | 2.74 | 12.41 | 2.54 | 2.43 | 0.89 | 3.96 | 0.0026 | 0.88 |
|  | 1500 | 48 | 9.96 | 3.05 | 12.23 | 2.50 | 2.27 | 0.65 | 3.89 | 0.0071 | 0.79 |
|  | 1200 | 48 | 9.98 | 2.92 | 12.32 | 2.45 | 2.34 | 0.77 | 3.91 | 0.0043 | 0.84 |
|  | 900 | 48 | 9.88 | 2.86 | 12.28 | 2.51 | 2.40 | 0.83 | 3.96 | 0.0034 | 0.85 |
|  | 600 | 48 | 10.03 | 3.24 | 12.60 | 2.57 | 2.57 | 0.87 | 4.27 | 0.0039 | 0.85 |
|  | 300 | 48 | 10.48 | 3.55 | 12.37 | 2.83 | 1.89 | 0.03 | 3.76 | 0.0470 | 0.52 |
|  | 1500 | 44 | 9.90 | 3.18 | 12.48 | 2.41 | 2.58 | 0.86 | 4.30 | 0.0043 | 0.84 |
|  | 1500 | 40 | 10.03 | 3.30 | 12.22 | 2.36 | 2.18 | 0.34 | 4.03 | 0.0216 | 0.65 |
|  | 1500 | 36 | 10.31 | 3.27 | 12.36 | 2.43 | 2.05 | 0.09 | 4.01 | 0.0404 | 0.55 |
|  | 1500 | 32 | 10.65 | 3.25 | 12.32 | 2.52 | 1.67 | -0.44 | 3.78 | 0.1158 | 0.35 |
|  | 300 | 44 | 10.40 | 3.68 | 12.53 | 2.79 | 2.13 | 0.13 | 4.12 | 0.0370 | 0.56 |
|  | 300 | 40 | 10.41 | 3.74 | 12.23 | 2.75 | 1.82 | -0.29 | 3.93 | 0.0882 | 0.40 |
|  | 300 | 36 | 10.69 | 3.72 | 12.32 | 2.86 | 1.63 | -0.62 | 3.89 | 0.1499 | 0.30 |
|  | 300 | 32 | 11.00 | 3.80 | 12.42 | 2.85 | 1.42 | -1.01 | 3.85 | 0.2421 | 0.21 |

Figure 12 suggests that with 300 stimulus presentations per participant, at least 44 participants would be required to establish a sex difference in p1-n1 latency. Only two of the studies described in table 1 meet these inclusion criteria (Welgampola & Colebatch, 2001; Shahnaz & David, 2021). Neither found a sex difference in p1-n1 latency. However, both studies are difficult to interpret on latency measures. One assessed 70 participants having an age range of 25–85 years (Welgampola & Colebatch, 2001) and did not report an age breakdown according to sex. Lack of report of an age breakdown according to sex introduces the possibility that latency changes due to age (Macambira et al., 2017) were unbalanced with regard to sex, making the study unsuitable for comparison of VEMP latencies based on sex. The other study was restricted to participants aged between 20–29 years, and found a sex difference in p1-n1 amplitude rather than p1-n1 latency (Shahnaz & David, 2021). However, this study barely met inclusion criteria (44 participants, with 400 or more presentations per participant). Its aim was to evaluate electrode montages rather than sex difference, and it contained a nested study of head turn versus head raise procedures which may have affected VEMP p1-n1 amplitude measurements (see section 2.5.2). Latency measurements were not reported in detail, with the absence of a sex difference implied and a comment that latencies were longer than in several other studies which used shorter duration stimuli. A study almost meeting criteria was that of Lee et al. (2008), who found n1 1.4 ms earlier in women than men (similar to 1.7 ms in the current study) and p1 trending earlier (unlike the current study, where p1 was prolonged in women compared to men). Lee et al. (2008) did not provide granular detail of sex matching with the wide age range (between 12 and 77 years) in their 97 participants, nor did they provide detail of the number of stimuli presented.

Figure 12 suggests that, with 300 or more presentations per participant, recruitment of more than about 44 participants is uneccesary. However, such a suggestion is not at all well supported by data. The most that could be inferred is that with exactly 48 participants, there seems little to be gained for measurements of VEMP p1-n1 latency by increasing presentation count from about 1000 to about 3000. At least two difficulties follow from such an inference. Firstly, the inference only applies when VEMP p1-n1 latency is the sole outcome measure. As described in sections 1 and 2, VEMP p1-n1 amplitude was also an outcome measure of interest. Assessment of VEMP p1-n1 amplitude requires a different statistical analysis (amplitude growth measurements, as reported in section 3.1) using a far greater amount of data than is necessary for VEMP latency assessment. From this perspective, the large presentation count available in the current study for analysis of VEMP latencies is not an indication of excess data collection, but is rather a side effect of the need to collect large quantities of data to accurately assess VEMP p1-n1 amplitude. Secondly, the current study presents no data upon which to model the effects of a participant count higher than 48. Thus, it may be that with a participant count higher than 48, studies can be adequately powered to detect a sex difference in VEMP p1-n1 latency even with fewer than 300 stimulus presentations per participant.

As regards this last-mentioned point, two further studies described in table 1 are pertinent. Brantberg et al. (2007) collected 192 VEMP responses from 1000 participants aged between 7 and 91 years. As described in section 2.5.5, the very high participant count in Brantberg et al. (Brantberg et al., 2007) will have reduced an inaccuracy in electrode positioning which acts as a confounder to VEMP latency measurements. The high participant count will also have reduced variation due to age. A granular age breakdown was not reported by Brantberg et al. (2007), but participants were consecutive visitors to a clinic over a period of nearly 3 years, with 625 women and 375 men. Brantberg et al. (2007) found no difference in n1 latency between women and men, however there was a significant age-sex interaction for women (p < 0.05) in which the p1 latency was not as prolonged with higher age for women as it was for men. The indication is therefore of p1-n1 inter peak latency becoming more prolonged for women than for men with age. Li et al. (2015) tested 186 participants aged between 26 and 92 years, but with a positively skewed age distribution (mean age was 73 years). Their age sampling was in contrast to the current study, in which all participants were aged between 16.6 and 21.1 years. Li et al. (2015) found p1 latency shortened by 0.39 ms in women compared to men (p = 0.005). The suggestion is of a longer p1-n1 inter peak latency for women versus men. There is again a contrast to the current study, which found a shorter p1-n1 inter peak latency for women versus men. Li et al. (2015) did not report n1 latency, making further comparison between their study and the current study difficult. However, it is worth noting (see section 3.3) that whereas Li et al. (2015) found p1 latency shortened by 0.39 ms in women compared to men, in the current study p1 latency was prolonged by 0.74 ms in women compared to men, albeit at the statistically insignificant p = 0.11 level.

For both sexes, p1-n1 latency has been found to increase across the lifespan. Meta-analysis between groups older or younger than 55 years showed that in the older group, p1 was prolonged by 1.2 ms (95% CI 0.2, 2.1) whilst n1 was prolonged by 2.8 ms (95% CI 0.3, 5.3) (Macambira et al., 2017). The sex-specific indication from available data is that between the ages of 17–21 years (current report) p1 is prolonged and n1 shortened in women compared to men. Across the lifespan, p1 becomes prolonged in both sexes, but moreso in men than in women (Brantberg et al., 2007). When the test group has a mean age of 73 years the p1 latency is prolonged in men compared to women (Li et al., 2015), which is the opposite of the situation at ages 17–21 years (current report). Caution is warranted with this interpretation, which is based on three cross-sectional studies which sometimes have incomplete data, e.g. around age/sex matching or n1 latency. However, it is presented here as the best summary of currently available data for sex difference across the lifespan in VEMP p1-n1 latency.

For the reasons described in section 2.5, evaluation of VEMP p1-n1 amplitude will require greater statistical power than evaluation of VEMP p1-n1 latency. Accordingly, only studies with more than the 44 participants necessary to evaluate sex differences in p1-n1 latency should be considered when assessing sex differences in p1-n1 amplitude. Of the prior studies meeting this criterion, five found no sex difference in p1-n1 amplitude (Brantberg et al., 2007; Li et al., 2015; Rosengren et al., 2011; Carnaúba et al., 2011; Blakley & Wong, 2015). These include the only two studies with substantially more than 100 participants (Brantberg et al., 2007; Li et al., 2015). Three studies did find a sex difference in p1-n1 amplitude (Welgampola & Colebatch, 2001; Lee et al., 2008; Shahnaz & David, 2021), with suggestion of an age effect, however all are subject to the confounders described in section 2.5. The current report minimised the confounders described in section 2.5, and found no sex difference in VEMP p1-n1 amplitude. Based on the current report and five prior studies (Brantberg et al., 2007; Li et al., 2015; Rosengren et al., 2011; Carnaúba et al., 2011; Blakley & Wong, 2015), the indication is of no sex difference in VEMP p1-n1 amplitude.

### 4.2 Possible causation unrelated to the vestibulo-collic reflex arc

Three possible causal explanations for the finding of sex difference in VEMP p1-n1 latency are described in this section. All are based on conduction outside the vestibular system. The first involves vibratory propagation from the transducer producing stimuli to hair cells in the vestibular system. The second involves the nature of the stimuli. The third involves electromagnetic conduction from the sternocleidomastoid muscle (SCM) to electrodes placed on the surface of the skin. In all of these explanations, the shortened p1-n1 latency in women compared to men would be a side effect of difference in non-vestibular physiology (e.g. of smaller head and neck volume in women than in men), rather than a difference between women and men in function of the vestibular system itself. Explanations involving conduction outside the vestibular system are supported by studies showing a smaller size for the temporal bone (Marcus et al., 2013), which includes the vestibular periphery, and for neck circumference (Vasavada et al., 2008; Machino et al., 2021) in women compared to men. Care is warranted though, because explanations within and without the vestibular system are not mutually exclusive. Thus, it is possible for both types of explanation to be true. It is moreover possible for both types of explanation to be true but to have different directions of fit (e.g. one prolongs latency, the other shortens latency), and it is possible that the effect from one type of explanation outweighs that from the other.

At first glance, an explanation based on non-vestibular conduction appears plausible. If head and neck size is smaller in women than men, the distance over which conduction occurs is also smaller. On the basis that propagation speeds for sound/vibration and electromagnetic radiation are constant, a shorter conduction time could be expected in women than in men. The consequence would be that both p1 and n1 latencies should be shorter for women than men. However, it is not obvious that latency difference between the p1 peak and the n1 trough should be shorter in women than in men, as was found in the current study. This is because if p1 and n1 latencies are equally affected by shorter conduction times, calculation of the p1-n1 inter-peak latency by subtraction should cancel out the effect of conduction time. Consider in this regard the study of Brantberg et al. (2007), described in section 4.1, which found that p1 latency became less prolonged in women than in men with increasing age, but that there was no age or sex effect on n1 latency. This describes two sets of asymmetries accompanying VEMP latency changes with increasing age: one is in p1 latency versus n1 latency, and another is in women compared to men. These asymmetries are not easily explicable using non-vestibular physiology. The reason for this is firstly that conduction effects would be expected to affect p1 and n1 equally, and secondly that men and women would be expected to be affected equally by the physiological changes which are associated with ageing (e.g. sarcopenia; Machino et al., 2021). There is moreover an effect size difficulty with an explanation involving vibratory propagation from bone conductor to vestibular mechanoreceptors. The mastoid bone placement for the bone conductor in the current study was within approximately 10 cm of vestibular mechanoreceptors. A propagation rate of 300 m/s for vibration through the head is likely in humans (Hotehama & Nakagawa, 2012). Thus, the direct propagation time for vibration from bone conductor to vestibular hair cells is less than 0.4 ms for either men or women. However, the sex difference in p1-n1 inter peak latency found in the current study was 2.4 ms. The propagation time from bone conductor to vestibular hair cells is in and of itself too short to explain the sex difference established in p1-n1 inter peak latency. Based on direct propagation of vibrational energy, and not on any other consideration, there is no plausible explanation for sex difference in VEMP p1-n1 latency.

A second prospective explanation for findings based on conduction outside the vestibular system concerns the nature of stimuli. The vestibulo-collic reflex arc begins with neural impulses along the VIII cranial nerve which are created when vestibular hair cells are deflected by vibrational energy. Stimulus type is thought to have an effect on the loci of hair cells which are deflected. For example, tests in guinea pigs have shown that air conducted stimuli preferentially activate the saccule (Curthoys et al., 2006) whereas body conducted stimulation is more evenly distributed between saccle and utricle (Curthoys et al., 2016). In humans, body-conducted vibration delivered to the mastoid appears to preferentially activate the utricle (Govender et al., 2015). Both saccule and utricle are otilith organs. Vestibular hair cells in a neuroepthilial layer are attached to crystalline structures in an otoconial layer, in such a way that the relative motions of the two layers will result in coordinated deflections of large numbers of vestibular hair cells, and hence coordinated variation to resting state action potential frequencies. For example, one account of the vestibular system’s response to vibration distinguishes between oscillatory modes in otoliths referred to as “accelerometer” and “seismometer” (Curthoys et al., 2019). When collecting VEMPs, two types of vibratory effect may alter the nature of hair cell deflection in the vestibular system. The first would depend on stimulus delivery (e.g. air- or body-conduction, including delivery point for body conduction). The second would involve oscillatory mode of otoliths. These effects would interact. They would moreover vary as a function of stimulus type (e.g. click or tone burst) and the vibratory modes within individual heads.

Vibratory modes within the skull are complex (Williams & Howell, 1990; Sohmer, 2017; Dobrev et al., 2017). Among several factors, vibratory modes depend upon head size. Thus, smaller head sizes in women than men could result in the oscillatory modes of otoliths differing between women and men for stimuli which are identical at the point of delivery. Suppose this biomechanical effect does occur, and that the differences in oscillatory modes alter firing rates of vestibular hair cells sufficiently to affect VEMP latency. Such a circumstance could account for the VEMP p1-n1 latency difference found between women and men in the current study. Cervical VEMPs have been recorded from click stimuli, and from tone bursts with duration of up to six milliseconds, delivered at a variety of locations (Rosengren et al., 2019). The nature of stimulation may crucially affect the response of the vestibular system. Support for this idea comes from the observation that delivery position and phase of body-conducted tone bursts affects VEMP amplitude and latency (Govender et al., 2016). Vestibular tuning has been found to depend on age (Janky & Shepard, 2009; Piker et al., 2013). It may also depend on sex. As described in section 4.1, participant and presentation counts are crucial for accurate appraisal of VEMP response. It may also be that stimulus type is a crucial factor when women and men are compared.

The third and final causal explanation based on conduction outside the vestibular system concerns the change in electromagnetic field created by the cessation in muscle spindle activity in the sternocleidomastoid muscle (SCM) during the vestibulo-collic reflex. Recordings from electrodes on the surface of the skin capture a compound muscle activation potential from many muscle spindles (Kane & Oware, 2012; Mallik & Weir, 2005). This summed activity is subject to volume conduction (Rutkove, 2007; Farina et al., 2002), a phenomenon in which the combined electromagnetic activity from many individual muscle spindles, along with electromagnetic propagation through non-excitable tissue (notably subcutaneous fat; Bartuzi et al., 2010) shapes electromagnetic potentials recorded on the skin surface. Thicknesses of the SCM and subcutaneous tissue have been measured with sonography (Chang et al., 2007), with the finding that raw amplitudes of VEMPs in adults correlated negatively with subcutaneous thickness. Despite this, the relationship with thickness of both subcutaneous tissue and SCM became statistically insignificant when VEMP amplitudes were corrected for pre-stimulus EMG background, as was the case in the current study. Thus, volume conduction appears at first glance unlikely to have been an appreciable factor affecting VEMP measurements.

However, there are two complications concerning electromagnetic conduction through the SCM. Both are related to neck length. Firstly, if electrodes were not positioned at the shortest possible distance from the electromagnetic source within the SCM, volume conduction will have increased.

Electrode placement for the current study was via a clinical palpation technique (BSA, 2012) which was consistent between women and men. The palpation technique aimed to position the electrode directly above the belly of the SCM. Locating the belly of the SCM via palpation will have become less accurate as a function of SCM size, which will have varied in turn as a function of neck length. Thus, a neck length difference between women and men could have led to a variation in electrode positioning which resulted in a sex difference in VEMP measurements, as was found in the current study.

The second complication concerns the nature of the VEMP. The VEMP has been described as a superposition of motor unit action potentials (Wit & Kingma, 2006; Lütkenhöner, 2019). Consider in this regard the proposition that the VEMP is a combination of several wave forms, including a standing wave and a travelling wave, which originate at a central motor point and propagate along the SCM in both directions (Rosengren et al., 2016). In such a scenario, a difference in SCM length between women and men could lead to a sex difference in standing waves and, through superposition with travelling waves, an alteration to VEMP p1-n1 latency as was found in the current study. Similar considerations will follow for other models of the VEMP involving superposition (Wit & Kingma, 2006; Lütkenhöner, 2019; Wei et al., 2013). Neck length once again emerges as an important factor.

In anthropometrical comparison, neck length in women was found to be no different from men between the sternum and tragus, but was 7 mm shorter between the C7 spinous process and tragus (Vasavada et al., 2008). However, these data were collected to assess sex difference pertaining to whiplash injury. As such, participants were closely paired on height and neck length, with an initial sample of 90 screened down to 28. Even then, neck variation could be substantial (e.g. 20 mm longer for the man than the woman in one pair). Another set of neck length data, from 88 participants, showed that the distance between the C7 spinous process and the external occipital protuberance was 9.5 mm shorter in women than men (Ahmed et al., 2020). The sex comparison was only made for participants with spondylosis, although comparison with non-spondlyosis controls showed that neck length was not a statistically significant predictor for spondylosis.

VEMPs have been compared between children and adults. The averaged findings were that in adults necks were 38 mm longer (mastoid tip to clavicle), p1 latency was 2.9 ms longer, n1 latency was 3.1 ms longer, and p1-n1 latency was 0.9 ms longer (Chang et al., 2007). Male and female necks were not measured in the current study. If male necks were longer than female necks then VEMP latencies were prolonged with increased neck length, similar to adults versus children. However, effect sizes differed: 1.7 ms for n1, and 2.4 ms for p1-n1 for men versus women in the current study, compared to 3.1 ms for n1 and 0.9 ms for p1-n1 for adults compared to children. Other relevant studies evaluated the effect of deliberately positioning active electrodes away from the belly of the SCM (Rosengren et al., 2016; Ashford et al., 2016). Rosengren et al. (2016) found that an active electrode position 50 mm from the belly of the SCM prolonged p1 and n1 absolute VEMP latencies by 3 ms. Needle electrode data were reported in integer milliseconds only, but surface electrode data for the same location were reported in sub-milliseconds, and show a prolongation of 2.3 ms in VEMP p1-n1 latency. These data are comparable to the 1.7 ms prolongation in VEMP n1 latency, and 2.4 ms prolongation in VEMP p1-n1 latency, found in the current study for men compared to women. Rosengren et al. (2016), tested three women and three men, which provided insufficient data for statistical comparison based on sex. However, the range of measurement in their study appears quite wide (e.g. the middle surface electrode n1 latency of 21.8 ms had a range of 17.4–28.0 ms, whilst a single motor unit needle electrode at the same location showed a latency of 12 ms with a range of 9– 21 ms). This raises the possibility that changing electrode positioning had differing effects on VEMP latency between women and men, and that such effects could have become apparent with a larger participant count.

Overall, there are insufficient data to draw a conclusion on whether differences between female and male necks could have contributed to the VEMP latency differences found in the current study. The indication is that volume conduction is unlikely to have been an appreciable factor. However, until more is understood of the precise nature by which the VEMP originates within and spreads throughout the SCM, the possibility of apparently small changes in neck dimensions having an appreciable effect on VEMP latency measurements cannot be ruled out.

### 4.3 Possible causation affecting the vestibulo-collic reflex arc

Another possible cause for the finding of a sex difference in VEMP p1-n1 latency may be a structural or functional difference between women and men which manifests in the vestibular reflex arc described in figure 1. Such explanations will consider activity in the vestibular periphery, the VIII and XI cranial nerves, the vestibular nucleus, the medial vestibulospinal tract and the sternocleidomastoid (SCM). Several of these areas were featured in the 2019 review of Smith, Agrawal & Darlington (2019). Relevant findings are summarised and extended here.

Quantities of type I and type II vestibular hair cells have not been found to differ significantly between men and women in any of the vestibular sensory organs (Merchant et al., 2000). Vestibular hair cells innervate, and for type II cells are innervated by, the bipolar neurons in Scarpa’s ganglion. Women have been found to have fewer Scarpa’s ganglion neurons than men (Velázquez-Villaseñor et al., 2000). Axons from Scarpa’s ganglion neurons comprise the vestibular portion of the VIII cranial nerve. Morphometric analysis of Scarpa’s ganglion axons has found no difference between women and men in average transverse area, and significantly fewer myelinated axons in women than in men (18,022 compared with 21,006; Moriyama et al., 2007). Other than the X cranial nerve, the vestibular portion of the VIII cranial nerve was the only one of 13 peripheral nerves to show a sex difference in a morphometric comparison (Moriyama et al., 2016). However, the XI cranial nerve, which innervates the sternocleidomastoid, was not included in the morphometric comparison, and on available data can only be evaluated for sex difference using a conduction study (Cleavenger et al., 2019). Latencies were assessed between the C7 spinous process and the trapezius muscle, in which the XI cranial nerve terminates after branching to the sternocleidomastoid. Latencies were up to 0.4 ms shorter in women than in men. This is too small a difference to account for the finding of the current study that VEMP p1-n1 latency was shorter in women than men by 2.4 ms.

Sex differences have been established in central components of the vestibular system. The vestibular system projects directly to cerebellar vermis (Hitier et al., 2014). Several studies have found that cerebellar vermis is larger in men than women (Raz et al., 1992, 1998, 2001; Tiemeier et al., 2010), although sometimes the opposite has been found (Luft et al., 1999) and some studies have found no group difference (Rhyu et al., 1999; Bernard et al., 2015; Metwally et al., 2021). Vestibular nuclei span the pons and medulla, which have been investigated in humans using diffusion tensor imaging (Bouhrara et al., 2021). Sex differences were found in the pons but not the medulla, via measures of fractional anisotropy, mean diffusivity and radial diffusivity. Measures of axial diffusivity, longitudinal and transverse relaxation rates, and myelin water fraction showed no sex difference. Ayyildiz et al. (2008) found the right medial vestibular nucleus (MVN) had a larger volume when comparing female to male rats. They also found a laterality difference, with male rats having more neurons in the left than the right MVN, and female rats having more neurons in the right than the left MVN.

In the brainstem generally, as in the cerebrum, the indication is that women have higher myelin content than men (Arshad et al., 2016; Bouhrara et al., 2020; Bouhrara et al., 2021). This is consistent with an interpretation in which proliferation of neuroglia and myelin proteins are regulated differently in women compared to men (Cerghet et al., 2006; Greer et al., 2004). Such a difference in regulation may be due to sex hormones (Marin-Husstege et al., 2004; Cerghet et al., 2009; Schumacher et al., 2012). Sex hormones were proposed by Ayyildiz et al. (2008) to play a crucial role in the sexual dimorphism they found in the MVN, with the contribution being primarily neurodevelopmental. Smith, Agrawal & Darlington (2019) reviewed a variety of studies demonstrating that the sex hormone oestrogen has an effect on brain areas considered to be important for the vestibular system, along with studies showing that chemicals known to be toxic to the vestibular system have a differing effect in female versus male animals. They also highlighted the shortage of studies appraising testosterone levels. A similar observation has been made by researchers in endocrinology (Singh Ospina et al., 2015), since androgens as well as oestrogens have been found to regulate critical biological and pathological processes in both males and females (Hammes & Levin, 2019). For example, injection of testosterone propionate has been found to reduce the effect of immune-mediated sensorineural hearing loss in female Wistar rats (Yeo et al., 2003). In male Long Evans Hooded rats, vestibular dysfunction caused by repeated mild traumatic brain injury was reduced in rats receiving testosterone, with the treatment improving vestibular neuronal survival by comparison to a sham rat group (Foecking et al., 2022). Testosterone has been found to mediate synaptic responses in MVN slices from male rats, depending on its conversion to estrogenic or androgenic metabolites (Grassi et al., 2010; Scarduzio et al., 2013), with vestibular synaptic transmission depending on estrous cycle in MVN slices from female rats (Pettorossi et al., 2011; Grassi et al., 2012). In humans, sex hormones have been linked to vestibular migraine, vertigo and Meniere’s disease (Tang et al., 2021; Park & Viirre, 2010; Seemungal et al., 2001). Indication overall is that sex hormones may affect VEMP measurements via an effect on myelination or synaptic response.

### 4.4 Use of BC stimuli to increase statistical power in VEMP testing

Use of AC stimuli in VEMP is crucial when ear specific information is required. This is due to intracranial conduction in which BC stimuli necessarily evoke a response from both ears, invalidating ear specific measures. This limitation of BC stimuli must be considered against the risk to hearing health with the high AC stimulus levels needed to evoke a VEMP response. Thus, for some vestibular research questions, the greater amount of data which can be safely collected with BC stimuli may be preferable to the ear specific measures available with AC stimuli.

Concerns around safe AC stimulus levels may even be understated, due to a tendency to underestimate sound pressure levels at the tympanic membrane when variation in ear canal size is considered. This follows from the use of a 2 cc coupler in standardised calibration procedures (e.g. ANSI S3.7). Correction for ear canals with volumes other than 2cc can be carried out on the basis that pressure is inversely proportional to volume (Boyle, 1662). From the definition of the dB scale, and using a 2 cc cavity as the reference, the necessary dB increase to dial SPL is 20 times the base 10 logarithm of the ratio of the 2cc coupler volume to the actual ear canal volume. Conveniently, it is not necessary to get involved in any actual pressure calculations. Rather, corrections can be made by addition or subtraction using log arithmetic, due to the corrections being transformations of the dB denominated pressure value relative to the 20 µPa standard. Thus, for an ear canal size of 1 cc, the correction is 20 log(2), or 6.02 dB. Correction for other ear canal volumes follows a similar arithmetic. For example, a 0.7 cc ear canal requires an increase to dial SPL of 20 log (2/0.7), or 9.1 dB. An ear canal size of 0.7 cc is not atypical for a woman, with ear canal size for both sexes having a 90% range between approximately 0.6 and 1.5 cc (Margolis & Heller, 1987). Thus, dial settings for dB (any scale) may be an appreciable underestimate when equipment is calibrated with a 2 cc coupler. This consideration is borne out in the study of Thomas et al. (2017), who measured ear canal size as 0.29 cc larger in a group of older compared to younger children using tympanometry. They found this corresponded to an approximate 3 dB increase in ear canal sound pressure for the younger children as recorded with a probe microphone. This compares to a predicted 2.9 dB increase using the formula just described.

In VEMP testing, 600 stimulus presentations per ear will typically provide sufficient data for clinical assessment (Rosengren et al., 2019). Based on an ear canal size of 2 cc and the calculations of Portnuff et al. (2017), 600 presentations of an AC 0-1-0 500 Hz tone burst over insert earphones would be within EU safe sound exposure limits at the typical 100 dB nHL stimulus levels used to elicit VEMP responses. However, if the ear canal size is revised to 1 cc, EU safe sound exposure limits would be exceeded. It follows that some clinics and research centres may unwittingly be exceeding safe sound limits during VEMP testing.

With BC stimuli, a total of 50,000 presentations of a 0-1-0 500 Hz tone burst would amount to less than 80% of EU safe sound exposure limits. Thus, there was effectively no restriction based on safe sound exposure limits for the amount of VEMP data which could be collected in single session using BC stimuli in the current study. As shown in figure 12, studies using fewer stimulus presentations could be underpowered. Previously published research using the same method (Gattie et al., 2021) showed a group difference in VEMP amplitude between participants who do and do not stutter. Indications in that study (see its figure 9, comparing a series of t-tests with small stimulus counts) are that the use of a greater number of stimulus presentations along with linear mixed effects regression modelling increased statistical power suffficiently to detect the group difference. Overall, the indication is that possibility of collecting greater quantities of data with BC stimuli can enable insights into the vestibular system which would be unavailable with AC stimuli due to the risk of exceeding safe sound exposure limits.

Following similar reasoning, use of BC stimuli may be preferred for initial VEMP assessment in the clinic. BC VEMP might be part of a test battery (i.e. alongside videonystagmography, head impulse test, rotary chair testing and so on) with AC VEMP testing following only in circumstances when a need for ear specific information is indicated.

## 5 Conclusion

VEMP p1-n1 latency was approximately 20% shorter in women than in men. There was no difference between women and men in VEMP p1-n1 amplitude. Comparison with prior studies indicated age is an important factor affecting VEMP latencies.

Candidate explanations for the finding include superposition of motor unit action potentials within the sternocleidomastoid, and influence of sex hormones. Sex hormones may affect myelination or synaptic response. Stimulus type and delivery may be important. The current study used a 500 Hz tone burst with rise/fall time of zero and a plateau of 2 ms, delivered to the mastoid bone using a bone conductor. Use of bone conduction will create vibratory modes within the skull which are not present using air conducted stimuli. This could lead to higher individuation, which may be sex dependent. The possibilities described are not mutually exclusive. Several sex difference may simultaneously occur. They may moreover have different directions of fit, with some effects outweighing others when an aggregate measure such as VEMP is taken.

The high AC stimulus levels required to evoke a VEMP response limit the quantities of data which can be collected in a single session. The current study and Gattie et al. (2021) have described a VEMP methodology in which use of BC stimuli enables collection of far greater quantities of data than would have been possible with AC data. Findings are of group differences which could have been missed if a limited delivery of AC stimuli had been used. Clinicians and research groups may find advantages in using BC stimuli for VEMP testing in the first instance, with AC stimuli used only when ear specific investigation is warranted.

